# How tp1, an indirect wing steering muscle, stabilizes *Drosophila’s* flight

**DOI:** 10.1101/2025.11.02.686144

**Authors:** Han Kheng Teoh, Debojyoti Biswas, Abby Leung, B. Kemper Ludlow, Sam Whitehead, Erica Ehrhard, Michael Dickinson, Tsevi Beatus, Noah J. Cowan, Itai Cohen

## Abstract

Flapping flight is inherently unstable, requiring *Drosophila* to fine-tune their wing motions on a milliseconds timescale. Pioneering studies have shown that the direct steering muscles, which attach to the wing hinge, are important for generating these rapid flight reflexes. Recent connectome data, however, reveal that indirect steering muscles or tension muscles, which alter the mechanics of the thorax, receive some of the same synaptic inputs from the sensory apparatus as the direct steering muscles. This discovery suggests that the indirect steering muscles may also be important for flight control. Here, we show that the indirect tergopleural muscles indeed contribute substantially to stabilization, particularly for large pitch perturbations. We find that for small perturbations (less than 1000 deg/s), flies modulate their wing stroke amplitude to generate lift-based corrective torques, a strategy that has been previously documented. For larger perturbations, however, we observe that *Drosophila* engage an additional wing degree of freedom—the wing pitch angle—to leverage additional lift and drag forces during the corrective maneuver. Quasi-steady aerodynamic simulations reveal that this strategy minimizes power consumption, anaolgous to how some mammals (including humans) adjust their steady-state gaits in a near energy-optimal manner. Using optogenetics and a control theory framework we demonstrate that the tergopleural muscle is activated by a proportional gain component of a nonlinear PI controller responsible for determining the wing pitch angle during large perturbations. A simplified torsional-spring model for the wing hinge captures the changes in the wing pitch dynamics observed during correction maneuvers by using the tergopleural muscle to adjust the rest angle of the wing. These findings provide a striking example of reflex strategy selection in time-critical behaviors and underscores the vital role of indirect steering muscles in flight stabilization.

## Introduction

Flapping flight, an extreme form of locomotion, demands continuous monitoring of flight stability and precise adjustment of wing motions on a millisecond time scale to maintain aerial control (Muijres et al., 2014; Taha et al., 2020; Chang and Wang, 2014). Insects like *Drosophila*, renowned for their rapid reflexes, demonstrate response times as short as 5 ms – some of the fastest in the animal kingdom (Beatus et al., 2015). Previous studies showed that these rapid correction strategies can be effectively described using proportional-integral (PI) controllers (Ristroph et al., 2010; Beatus et al., 2015; Whitehead et al., 2015, 2022), which is a powerful framework for modeling sensorimotor feedback control in various animals (Mussells Pires et al., 2024; Peterka, 2018; Stöckl et al., 2017; Sutton et al., 2016; Sponberg et al., 2015; Fuller et al., 2014; Schnell et al., 2014; Dyhr et al., 2013; Sefati et al., 2013; Roth et al., 2012; Lee et al., 2008; Lockhart and Ting, 2007; Madhav and Cowan, 2020; Li et al., 2025). For example, when flies are perturbed about the pitch axis, they adjust the forward sweep angle of the wings, which generates appropriate corrective torques that can counteract the aerial disturbances (Whitehead et al., 2015). Recent work has shown that a PI controller framework that incorporates the fly’s body angular velocity (P) and the total angular displacement (I) with appropriate coefficients can predict this change in forward sweep angle. Similar PI frameworks have been used to predict the wing modulations used to counteract perturbations along the yaw (Ristroph et al., 2010) and roll (Beatus et al., 2015) axes. In addition to being predictive of the wing modulations, these frameworks provide a foothold for understanding the neuromuscular implementations of flight control.

To maintain flight while executing this rapid and robust aerial control, the *Drosophila* flight motor system is divided into large asynchronous muscles that generate the high-frequency wing strokes and small synchronous wing steering muscles that implement rapid, fine adjustments to the wing motions. The large asynchronous muscles – the dorsal-longitudinal and dorsal-ventral muscles – form an antagonistic oscillatory system that produces wing beat frequencies of 200-250 Hz through reciprocal stretch activation. The small synchronous muscles can be classified as direct or indirect steering muscles based on their attachment within the wing hinge. Direct steering muscles attach to sclerites, the skeletal elements of the hinge, while indirect muscles influence wing motion via mechanical alteration of the thorax (Dickinson and Tu, 1997; Lindsay et al., 2017; Trimarchi and Schneiderman, 1994; Melis et al., 2024). The direct wing steering muscles are anatomically divided into four major groups based on the sclerites they insert on: the basalars (b1, b2, and b3), the first axillaries (i1 and i2), the third axillaries (iii1, iii3, and iii4), and the fourth axillaries (hg1, hg2, hg3, and hg4). In the context of flight, a large amount of effort has focused on understanding the role of these direct steering muscles. For example, experiments in tethered flies have determined via Ca^2+^ imaging the activity of the steering muscles in response to various visual stimuli (Lindsay et al., 2017; Melis et al., 2024). Such experiments have shed light on which muscles may be recruited to execute various aerial maneuvers associated with the pitch, roll, and yaw degrees of freedom. In addition, since many of these direct steering muscles are innervated by a single motoneuron, studies have been able to use optogenetic manipulation to determine the functional role of the muscle being activated in free-flight maneuvers. Notably, it was recently shown that the basalar motor units b1 and b2 respectively modulate the I and P elements of the PI controller for the pitch degree of freedom (Whitehead et al., 2022). As these discoveries highlight, this active area of study is rapidly advancing our understanding of the neuromuscular organization that distributes wing actuation across this flight motor system to generate such precise maneuvering and control in *Drosophila*.

Surprisingly, however, the role of the indirect wing steering muscles – tergopleurals (tp), pleurosternals (ps), and tergotrochanter (tt) – in flight control remains largely unexplored. Based on past morphological studies, the indirect wing steering muscles’ influence on wing motion is thought to arise from alterations to the mechanics of the thorax. Recent research has started to elucidate their crucial role in courtship. For example, the indirect wing steering muscles were shown to be important in modulating the wing motions responsible for ‘pulse’ (200-280 Hz) and ‘sine’ (160 Hz) songs (O’Sullivan et al., 2018; Ehrhardt et al., 2023). Both courtship song modes require sustained wing vibrations, similar to the wing flapping frequency during flight (200-250 Hz). These findings show that the indirect wing steering muscles are capable of patterning the wing profile within a wing beat cycle. Moreover, previous studies have shown that flies with chronically silenced tp muscles have their wing postures modified and their flight abilities significantly impaired (O’Sullivan et al., 2018). Finally, premotor synaptic connectivity data (Lesser et al., 2024) shows that many of these muscles receive haltere sensory inputs that are shared with the direct steering muscles. Such results indicate that despite the lack of attention paid to the indirect steering muscles, they may, in fact, be important for flight.

Here, we examine the role of the tergopleural or tp muscles, in corrective flight maneuvers. In most insect species, including *Drosophila*, the tp muscles consist of two primary components, tp1 and tp2, while in *Tabanids* (horse flies) and *Eristalis* (hoverflies), these muscles are fused into a single component (Miyan and Ewing, 1985). The tp muscles are speculated to modify the wing angle of attack and wing beat frequency by modulating thoracic stiffness (Dickinson and Tu, 1997). Unlike other wing steering muscles, which are each innervated by a single motoneuron, the tp1 and tp2 muscles receive innervation from their respective tp1 and tp2 motoneurons, as well as an additional neuron known as the tpn motoneuron that stochastically innervates both muscles (O’Sullivan et al., 2018; Ehrhardt et al., 2023; Azevedo et al., 2024). We discover that these muscles are important for implementing one of the reflex strategies for stabilizing pitch, and that this tp-mediated reflex is engaged during large amplitude pitch perturbations.

## Results

### The Tergopleural muscle alters the pitch stabilization reflex

To gain insight into the potential influence of the tp muscles (Fig. 1a) on flight reflexes, we determine whether they share any synaptic haltere sensory inputs with the b1 and b2 direct wing steering muscles, which have been previously studied in the context of pitch stabilization. To determine these shared synaptic inputs, we utilized the recently published premotor synaptic connectivity data (Lesser et al., 2024) reconstructed from the (Female Adult Nerve Cord) FANC dataset (Azevedo et al., 2024). We searched for premotor neurons that project from the halteres – the primary sensor used for rapid flight control (Mohren et al., 2019; Nalbach and Hengstenberg, 1994; Pringle, 1948; Dickerson et al., 2019) – and innervate the b1, b2, and tp motoneurons (Supp. Fig. S1). We found five premotor neurons that not only synapse onto the b1 or b2 motoneurons but also innervate the tp1 motoneuron (Fig. 1bi, Supp. Fig. S2). In particular, three premotor neurons innervate the b1, b2, and tp1 motoneurons; one premotor neuron innervates the b1 and tp1 motoneurons; and another premotor neuron innervates the b2 and tp1 motoneurons. The fact that the tp1 motoneuron shares premotor inputs with the b1 and b2 motoneurons suggests the tp1 muscle may play a similar functional role as the basalar muscles in the pitch stabilization reflex. Interestingly, we did not find any premotor neurons that synapse onto either b1 or b2 and also innervate either the tp2 or tpn motoneurons. We hypothesize that if the tp2/tpn motoneurons were to be involved in the pitch stabilization reflex, their recruitment would be indirect, perhaps based on local connectivity among the tp motoneurons. The remainder of our analysis focuses on the tp1 motoneuron’s role in stabilizing pitch (See SI for analyses of the tp2, and tpn motoneurons’ influence on flight control. We find data trends similar to those we report for tp1 but with higher variability, which made them more difficult to analyze for significance). To investigate the tp1 muscle’s impact on free flight, we take advantage of recently developed transgenic lines (Ehrhardt et al., 2023; O’Sullivan et al., 2018) that target the tp1 motoneuron (Fig. 1bii), which innervates the tp muscle (Fig. 1biii).

**Figure 1.**
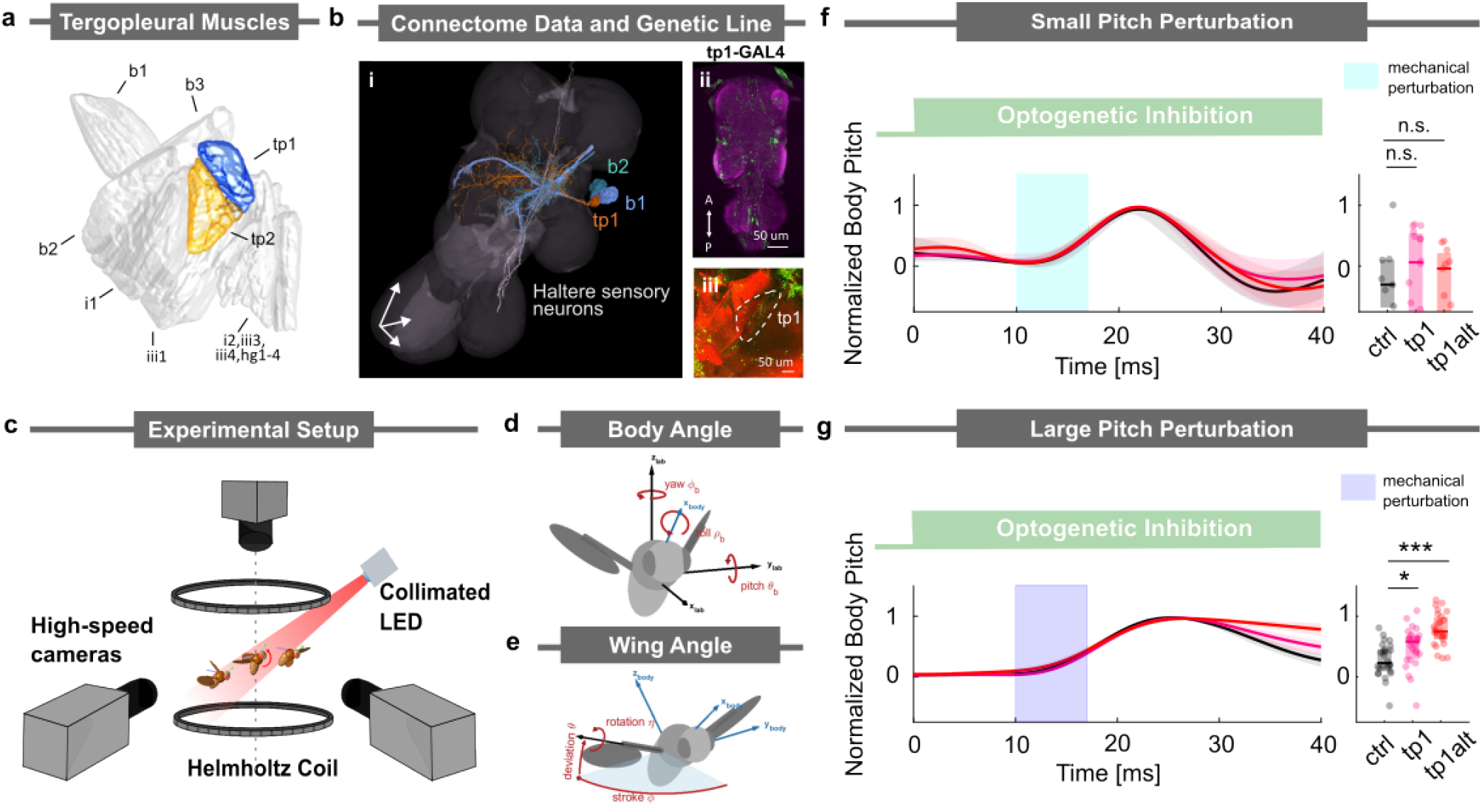
The tergopleural muscle, tp1, is involved in stabilizing flight during a large pitch perturbation. (a) Direct and tension wing steering muscles of *Drosophila*, with the tergopleural muscles (tp) highlighted in orange and blue. Data taken from (Lindsay et al., 2017). (b) i) Identified haltere premotor neurons (white) within FANC dataset that simultaneously innervate b1(blue),b2(green), and tp1(orange) motoneurons, ii) Maximum intensity projections of the fly ventral nerve cord (VNC) expressing the split line tp1-GAL4. Scale bars, 50 *µ*m. iii) Phalloidin-stained thoracic hemisection from a tp1-GAL4 > CsChrimson fly showing wing musculature, with both phalloidin (red) and GFP (green) expression innervating the tp1 muscle (outlined with white dashed lines). Scale bars, 50 *µ*m. (c) Schematic of the experimental apparatus used to deliver optogenetic and/or mechanical perturbations to freely flying *Drosophila*s. The magnetic field from Helmholtz coils is used to perturb flies with a magnetic pin glued to their notum. These magnetic perturbation events are filmed at 8000 frames/s. (d and e) Definitions of the body and wing Euler angles used to describe flight kinematics. (f, g) Body orientation along the pitch axis during correction maneuvers in the (f) small and (g) large perturbation regimes. Left: Average time traces of normalized body pitch angle. The green bar indicates optogenetic inhibition; the cyan/blue bar indicates mechanical perturbation. Right: Normalized body pitch angle at 40 ms. For the large perturbation condition, a Kruskal–Wallis test revealed a significant effect across groups (*H* = 28.7753, *p <* .001, *η*^2^ = 0.3523). Post hoc Dunnett’s tests showed a significant difference between control and tp1 (*p* = .0224), and a highly significant difference between control and tp1-alt (*p <* .001). For the small perturbation condition, no significant group differences were observed (*H* = 1.08, *p* = .584, *η*^2^ = 0). Sample sizes for small perturbation: control *n* = 9, tp1 *n* = 16, tp1-alt *n* = 11; for large perturbation: control *n* = 26, tp1 *n* = 27, tp1-alt *n* = 26.

To examine the role of the tp1 muscle in the pitch stabilization reflex, we optogenetically silenced the tp1 motoneuron while applying a pitch perturbation in free flight. We targeted the tp1 motoneuron using the split-GAL4 driver line (Ehrhardt et al., 2023), SS41052-GAL4 (Fig. 1bii,biii) to drive expression of GtACR1 (inhibition). In our free-flight pitch perturbation behavioral assay (Fig. 1c), we optogenetically silenced the tp motoneurons by delivering a 45 ms light pulse. 10 ms after the LED onset, we applied a 7 ms magnetic perturbation. We captured the fly’s pitch correction maneuver for various pitch deflection amplitudes using high-speed videography and extracted body and wing kinematics (Fig. 1d,e; Materials and Methods) (Ristroph et al., 2009; Whitehead et al., 2015). We discovered that for small pitch perturbations (deflection amplitude less than 10°), both the control and tp1-silenced flies were able to return to their original orientation within ~30 ms (Fig. 1f). However, when presented with a large pitch perturbation (deflection amplitude greater than 20°), the control flies were able to return to their original orientation in ~40 ms, which is slightly longer due to the larger perturbation amplitude. Here, the tp1-silenced flies recovered more slowly than the control group (pink curve in Fig. 1g). To control for any potential influence in our experiments from off-target expressions, we repeated our experiments using an alternative driver line, tp1-SG (tp1 alt), which was generated from a different parental lineage (O’Sullivan et al., 2018) (see Materials and Methods). We observed a degradation in correction performance during large pitch perturbations in flies from both driver lines (red curve in Fig. 1g). These observations led us to hypothesize that flies adopt a distinct pitch stabilization strategy in the large perturbation regime and that silencing of the tp1 muscle affects the implementation of this strategy.

### Fruit flies employ an additional reflexive strategy to correct for large pitch perturbations

To assess whether fruit flies employ a distinct pitch stabilization strategy in response to large perturbations, we first characterize the wing kinematic responses across both small and large perturbation regimes (see schematics in Fig. 2). We found that when a fruit fly is given a small-amplitude pitch-up perturbation of ~10°, the fly exhibits a reduction of ~20° in its front stroke angle (Fig. 2a). This reduction in the front stroke angle limits the forward extent of wing motion, thereby decreasing the anteriorly generated aerodynamic torque that leads to a net corrective moment that pitches the fly forward. A consistent decrease in front stroke angle at the wingbeat corresponding to the maximum corrective response was observed across all 29 flies subjected to perturbations of less than 10°(Fig. 2b). In contrast, comparisons of wing pitch angle trajectories before and during the perturbation revealed no significant differences (Fig. 2c–d), indicating that corrective responses in this regime are primarily mediated through stroke angle modulation. These findings are consistent with previously identified pitch correction mechanisms in fruit flies (Whitehead et al., 2015).

**Figure 2.**
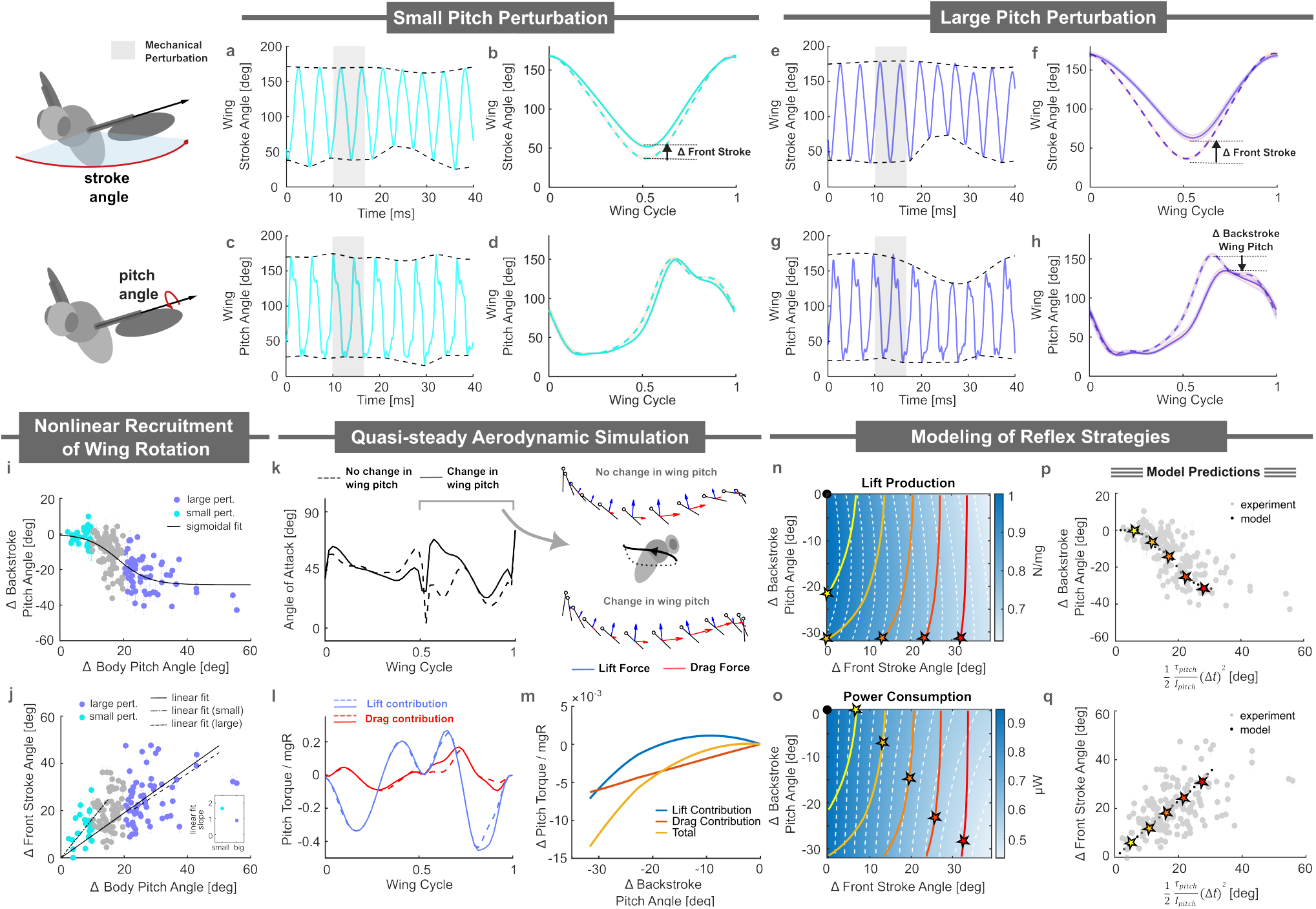
*Drosophila* use two distinct modes to stabilize pitch. (a, c) Wing stroke and wing pitch angles of a fly encountering a small pitch perturbation. The fly modulates its front stroke angle in response to body pitch instability. (b, d) Averaged wing stroke and wing pitch profiles (n=29 flies) for pre-maneuver motion (dashed line) and the largest corrective response (solid line). (e, g) Wing stroke and wing pitch angles of a fly encountering a large pitch perturbation. The fly modulates both the front stroke angle and backstroke wing pitch angle in response to the instability. (f, h) Averaged wing stroke and wing pitch profiles (n=61 flies) for pre-maneuver motion (dashed line) and the largest corrective response (solid line). (i) Δ Backstroke wing pitch angle response as a function of body deflection amplitude. The black curve represents a sigmoidal fit. Cyan points indicate experimental data for pitch deflection less than 10°; purple points indicate data for pitch deflection greater than 20°. (j) Change in front stroke wing angle (Δ front stroke) as a function of body pitch deflection amplitude. Solid black lines corresponds to linear fit to full dataset and dashed black lines are associated with linear fits across small and large perturbation regimes. The inset shows the slope of the linear fits in each regime for comparison. (k) Angle of attack as a function of wing cycle for two wing profiles representing small (dashed) and large (solid) perturbation responses. Differences in the angle of attack during the backstroke are illustrated by ball-and-stick diagrams, with red and blue arrows denoting drag and lift forces, respectively. (l) Drag (red) and lift (blue) contribution to corrective pitch torque normalized by the fly’s mass, wing span, and gravitational acceleration *mgR* for two wing profiles representing small and large perturbation responses. The dashed line indicates no backstroke wing pitch modulation; the solid line represents a 20°backstroke wing pitch modulation. (m) Contribution of drag and lift forces to corrective torque as the backstroke wing pitch angle varies. (n) Lift force generated as front stroke and backstroke wing pitch angles vary from stable flight (black point). The yellow to red curves show combinations that produce the same corrective torques. The star marks the optimal wing kinematics based on a lift maximization strategy. (o) Similar to (n), but examining power consumption. The star marks the optimal wing kinematics based on a power minimization strategy. (p, q) Backstroke wing pitch and front stroke responses generated using the power minimization strategy. Grey points represent experimental data from (i) and (j) for comparison.

When facing a large-amplitude pitch-up perturbation, however, flies implemented an additional corrective mechanism. For example, when a fly was subjected to a ~37° pitch perturbation, the fly not only reduced its front stroke angle by ~40°—similar to the small-perturbation response (Fig. 2e)—but also exhibited a ~40° decrease in wing pitch angle (Fig. 2g), thereby increasing the angle of attack during the backstroke. This dual response was consistently observed across all 61 flies experiencing large pitch perturbations (Fig. 2f,h). To characterize the transition between regimes, we plotted the change in backstroke wing pitch angle (Δ Backstroke Wing Pitch) against body pitch deflection (Δ Body Angle), which we found is reasonably well fit by a sigmoidal relationship that captures the expected asymptotic behviors at high and low perturbation amplitudes (Fig. 2i; Sigmoidal fit AIC : 686< Linear fit AIC: 694 see Method G). These data suggests that flies began to engage wing pitch modulation for perturbations greater than ~10°. Notably, changes in front stroke angle (Δ Front Stroke) were broadly similar across both perturbation regimes (Fig. 2j), though the stroke response had a steeper slope for data restricted to the small perturbation regime. Together, these results revealed a transition from a low-amplitude response strategy that relied predominantly on front-stroke modulation to a high-amplitude regime that engaged wing pitch modulation.

To understand how backstroke wing pitch modulation contributed to overall pitch corrective torque, we performed quasi-steady aerodynamic calculations (Materials and Methods) comparing the maximum torques for pitch perturbations with and without the added wing pitch response. In these simulations, we mixed and matched the wing kinematics for stroke, deviation, and wing pitch degrees of freedom. Specifically, we compared the pitch torques generated by wing kinematics when responding to a large pitch perturbation with those generated using the same stroke and deviation angles but retaining the wing pitch trajectories from pre-perturbation kinematics. Overall, the wing pitch angle contributes an additional ≈25% corrective pitch torque. We examined the differences in angle of attack and aerodynamic forces between these two sets of wing kinematics (Figs.2k–l). We found that lowering the wing pitch angle in the large perturbation regime increased the angle of attack during the backstroke. To emphasize this effect, we included an accompanying illustration featuring a “ball-and-stick” diagram, representing the wing’s leading edge and chord, along with visualizations of the corresponding lift (blue arrows) and drag (red arrows) forces during the backstroke phase. This diagram highlights larger drag forces (shown as more prominent red arrows) and relatively unchanged lift forces (shown as comparable blue arrows) associated with wing pitch angle modulation. Further analysis of the contributions from drag- and lift-based torques across a range of wing pitch angles revealed distinct trends: drag forces contributed linearly to the corrective torques, whereas lift forces exhibited a quadratic relationship with wing pitch angle (Fig. 2m). In prior investigations, it was shown that by operating near an angle of attack of 45°, where lift forces are maximal, flies are able to modify drag forces (linear dependence on angle of attack) while keeping lift forces (quadratic dependence on angle of attack) relatively unchanged. Here, we find that the wing angle of attack is sufficiently altered that both lift and drag forces contribute significantly to the corrective torque. Collectively, these results indicate that the correction strategy under large perturbations also involves wing pitch modulation.

These results raised an important question: why did fruit flies implement this strategy during these large perturbations? In principle, a fruit fly could have achieved the same corrective torque in this regime by simply increasing the modulation of its front stroke angle. However, such a modulation would have altered the total lift generated, which is essential for keeping aloft. This observation suggests that the flies might have adopted a strategy aimed at maximizing lift. Alternatively, the shift in strategy may have been driven by energetic considerations, such as minimizing power consumption. To explore these possibilities further, we developed a simulation-based model (see Supplementary Information) that allowed us to analyze the interplay between lift, power, and torque production. The model parameterizes wing strokes using variations in the backstroke pitch angle (Δ Backstroke Pitch Angle) and front stroke angle modulation (Δ Front Stroke Angle).

To investigate whether the fly was attempting to maximize its lift, we created a lift production heat map and overlaid equal-lift contours (dashed white lines and color bar in Fig. 2n). Superimposed on this heat map are equal pitch torque contours (solid colored lines, ranging from yellow to red, representing dimensionless torques of −0.04, −0.10, −0.16, −0.22, −0.28, and −0.34). The black point represents a stably hovering fly, experiencing zero aerodynamic torques and producing a total lift force sufficient to support its weight. The stars indicates the optimal adjustments to the Δ Backstroke Pitch Angle and Δ Front Stroke Angle needed to generate a specific torque while maximizing lift. Our analysis revealed that implementing this strategy would have entailed initially adjusting the backstroke pitch angle to achieve the desired dimensionless pitch torque. Once the fly reached its wing pitch rotational limit, here set at 40°, it would have resorted to changing its stroke angle degree of freedom (see also Supplementary Fig. S6). This strategy, however, differed from what we observed in the experiments (Fig. 2i), thereby ruling out the hypothesis that the fly’s behavior was driven by lift maximization.

Next, we plotted a power consumption heat map along with equal-power contours (dashed white lines and color bar in Fig. 2o) to investigate whether the fly aimed to minimize its energy expenditure. Once again, we overlaid equal pitch torque isolines onto the heat map (white dashed lines). The black point represents a stably hovering fly, experiencing zero aerodynamic torques and producing a total lift force sufficient to support its weight. The stars indicates the optimal adjustments to the Δ Backstroke Pitch Angle and Δ Front Stroke Angle required to generate a specific torque while minimizing power consumption. Under the power minimization hypothesis, we found that the front stroke angle was initially adjusted to generate small dimensionless pitch torques, as this required the least power. For larger dimensionless pitch torques, both the backstroke pitch angle and the front stroke angle were utilized, as this combination minimized power expenditure along the pitch torque isolines.

To better illustrate this trend, we simulated a fly correcting for a pitch perturbation of a given magnitude and extracted the Δ Backstroke Pitch Angle and Δ Front Stroke Angle under the constraint of power minimization. Specifically, we calculated the pitching torque for each body pitch angle, *τ*_pitch_, required to return the fly to its original orientation within a time scale of two wing beats (Δ*t*). Using the power consumption landscape from Fig. 2o, we identified the optimal adjustments to the Δ Backstroke Pitch Angle and Δ Front Stroke Angle and plotted them in Fig. 2p and Fig. 2q, respectively. These model-derived plots closely resemble their experimental counterparts (Fig. 2i,j). Notably, we observed a sigmoidal change in the Δ Backstroke Pitch Angle and a linear increase in the Δ Front Stroke Angle. The resulting picture suggests that the fly introduced the drag-based correction mechanism to minimize power consumption during large perturbations.

### The tergopleural muscle affects the fly’s ability to implement the large perturbation stabilization reflex

These results, in conjunction with the data from Figs. 1f,g, suggested that the tp1 muscle played a critical role in generating the wing stroke changes for the minimal power consumption correction strategy in the large perturbation regime. To further investigate the role of the tp1 muscle in implementing these strategies, we analyzed the wing kinematic changes resulting from optogenetic inhibition and activation of the tp1 motoneuron in both small and large perturbation regimes. We focused our analysis on the normalized changes in the backstroke wing pitch response and front stroke angle response, as they were the key drivers for generating the corrective torques for both strategies.

We show the results of optogenetic inhibition on these wing responses in Fig. 3a–d. For small perturbations (Fig. 3a,b), optogenetic inhibition of the tp1 motoneuron (red) did not significantly alter the wing pitch and wing stroke angles from those observed in control (black) experiments. On the other hand, we found that for large perturbations (Fig. 3c,d), the backstroke wing pitch response of the tp1-inhibited flies (red) was significantly smaller than that observed in the control experiments (black). Once again, we did not observe any statistically significant differences between the peak responses in the silenced and control groups for the maximum change in normalized front stroke angle. We did, however, observe a consistent decrease in the population average response over the entire correction maneuver. These data indicated that during normal correction maneuvers, the tp1 muscle did not alter the wing kinematics in the small perturbation regime. In the large perturbation regime, this muscle primarily controlled changes to the backstroke wing pitch angle and may have modulated the wing stroke angle as well. These wing stroke changes resulted in a decrease in the proportion of tp1-inhibited flies that corrected more than 50% of body deflection amplitude at 40 ms (Fig. 3e) compared to the control group.

**Figure 3.**
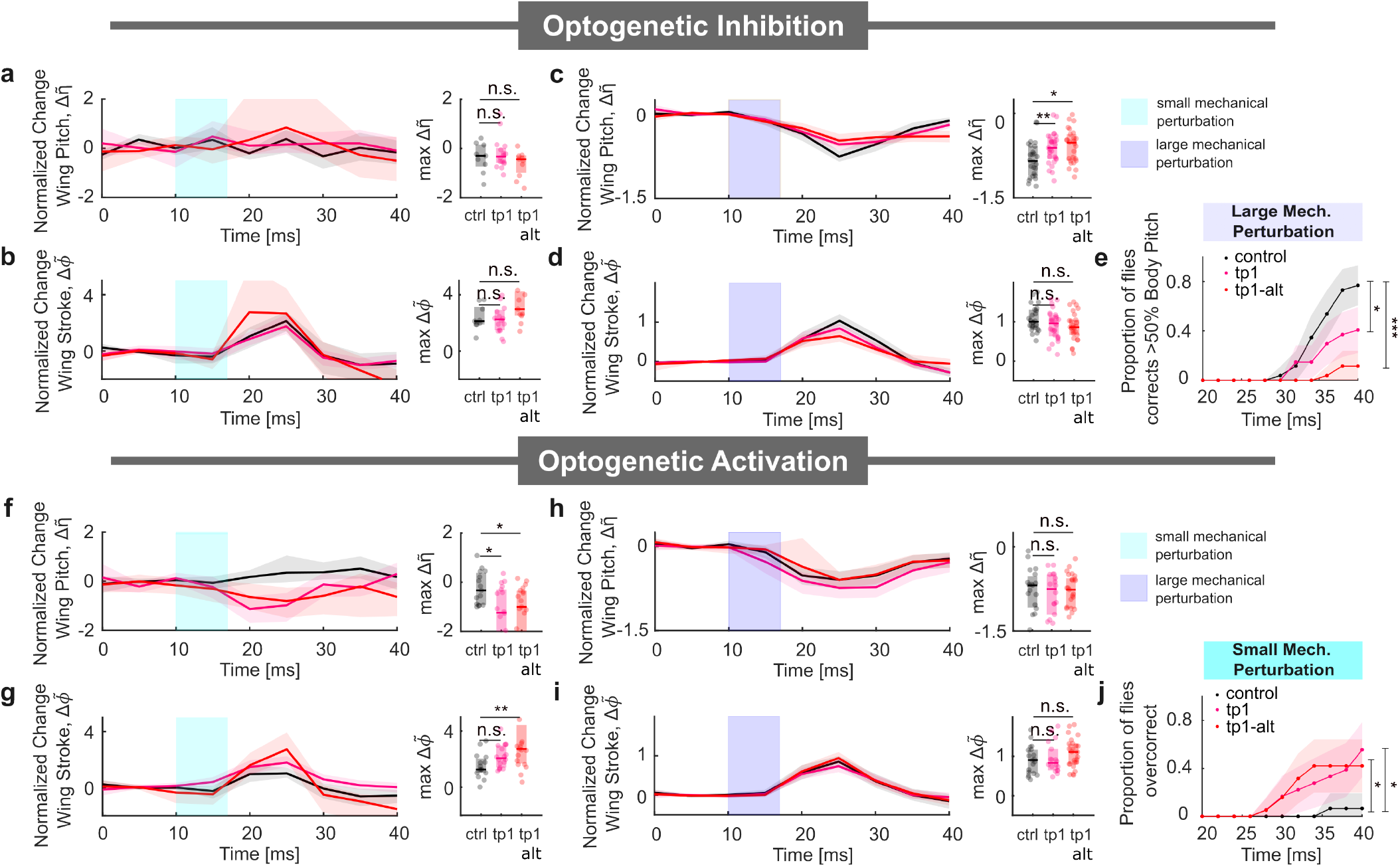
The tp1 muscle is involved in implementing the wing pitch response. (a, b) Left: Normalized change in backstroke wing pitch 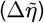 and front stroke 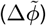 response for control and tp1-inhibited flies during small pitch perturbations. Right: Maximum change in normalized wing pitch and wing stroke responses. (c, d) Left: Backstroke wing pitch 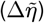 and front stroke 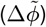 response for control and tp1-inhibited flies during large pitch perturbations. Right: Maximum change in normalized wing pitch and wing stroke responses. (e) Proportion of flies correcting more than 50% of the control flies’ body deflection amplitude at 40 ms. (f, g) Left: Normalized change in backstroke wing pitch 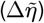 and front stroke 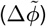 response for control and tp1-activated flies during small pitch perturbations. Right: Maximum change in normalized wing pitch and wing stroke responses. (h, i) Left: Backstroke wing pitch 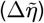 and front stroke 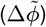 response for control and tp1-activated flies during large pitch perturbations. Right: Maximum change in normalized wing pitch and wing stroke responses. (j) Proportion of flies over-correcting more than 10% of the control flies’ body deflection amplitude at 40 ms. Statistical significance for (a)–(d) and (f)–(i) was determined via the Kruskal–Wallis test with Dunn’s post hoc multiple comparisons. Statistical significance for (e) and (j) was determined via the proportional test with Bonferroni correction (\*\*\**p <* .001; \*\**p <* .01; \**p <* .05). Full statistical results are provided in Supplementary Table 2. Sample sizes for optogenetic inhibition with small perturbation: control *n* = 12, tp1 *n* = 24, tp1-alt *n* = 15; with large perturbation: control *n* = 15, tp1 *n* = 26, tp1-alt *n* = 19. Sample sizes for optogenetic activation with small perturbation : control *n* = 16, tp1 *n* = 17, tp1-alt *n* = 11; with large perturbation : control *n* = 19, tp1 *n* = 16, tp1-alt *n* = 11.

Next, we investigated the effects of optogenetic activation on the wing responses (Figs. 3f–i). We found that in the small perturbation regime (Fig. 3f,g), optogenetic activation of the tp1 motoneuron resulted in significant changes to the maximum response of the normalized backstroke wing pitch angle (Fig. 3f). For the maximum change in normalized front stroke angle, we observed slight increases during activation. For the tp1-GAL4 lines, we did not observe any significant differences from the control group for the peak response, though the wing stroke angle was consistently higher over the course of the entire correction maneuver (Fig. 3g). For the tp1-SG line, we observed a higher, and statistically significant, maximum change in the normalized front stroke angle.

In the large perturbation regime (Fig. 3h,i), we observed no significant changes in either the normalized backstroke wing pitch or front stroke angle responses. The optogenetic activation of the tp1 motoneuron in the small perturbation regime indicated that additional pitch torques were generated via changes in the wing pitch angle and the wing stroke angle, which could lead to an over-correction. To test this conjecture, we counted the proportion of flies within each group that corrected beyond the 10th percentile of the control group body pitch angle at 40 ms (Fig. 3j). We observed a significantly higher proportion of tp1 motoneuron-activated flies that over-corrected (also see Supp. Fig. S7a). The data for the large pitch perturbations were consistent with the idea that the tp1 muscle was already active in the large perturbation regime; since the tp1 muscle would already have been active in this regime, optogenetic activation did not affect the wing kinematics. Consequently, we also did not observe a significant difference in the normalized body pitch angle response when the tp1 motoneuron was activated (Supp. Fig. S7b).

### Tergopleural muscle controls the rest angle parameter in the torsional spring model for the wing hinge

The wing hinge of the fly is a complicated joint that uses coordinated actions of the sclerite processes at the wing base, as modulated by various wing steering muscles, to generate the wing motions. Prior studies, based largely on static morphological observations across a range of dipteran species (Boettiger and Furshpan, 1952; Pringle, 1957; Wisser and Nachtigall, 1984; Miyan and Ewing, 1985; Ennos, 1987; Dickinson and Tu, 1997; Walker et al., 2014; Deora et al., 2017), have provided a mechanistic perspective on how wing steering muscles influence wing motion. For example, the tergopleural (tp) muscles were shown to produce small relative movements between the scutum and the pleural wing process, leading to a rotation of the first axillary sclerite. This rotation was hypothesized to determine the positioning of the radial stop on the pleural wing process, and consequently to affect the angle of attack of the wing (Miyan and Ewing, 1985).

Given the complexity of the wing hinge, here, we simplify our analysis by examining the effects of the tp muscles within a simple torsional spring framework. While this model has not been directly tested due to the challenges of measuring wing hinge stiffness and damping during free flight, it nevertheless provides a useful conceptual simplification that captures several aspects of wing stroke dynamics. For example, previous studies have shown that the dynamics of the wing pitch angle can be effectively described using a damped torsional spring model (Bergou et al., 2007; Beatus and Cohen, 2015). This approach captures the essential features of wing pitch dynamics by treating the wing hinge as a torsional spring resisting wing rotations induced by aerodynamic and gravitational torques (Fig. 4a). The torsional spring torque, *τ*_spring_, is expressed as 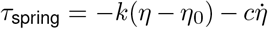, where *k* is the torsional spring stiffness, *c* is the damping factor, and *η*_0_ is the rest angle. Although this model is highly simplified and does not fully capture the detailed mechanics of the wing hinge, it offers a more intuitive perspective on how wing rotation might be modulated by the muscle activity through changes in the spring parameters. Here, we applied the torsional spring model (Bergou et al., 2007; Beatus and Cohen, 2015) to investigate the changes in spring parameters underlying the observed wing pitch kinematics in the control group. Using a similar approach outlined in a previous study (Beatus and Cohen, 2015), we utilized the body and wing kinematics (excluding wing pitch) for each wingbeat along with a set of optimized spring parameters to best replicate the experimental wing pitch dynamics (Fig. 4b-c). We then repeated this procedure on a stroke-by-stroke basis to examine changes in the spring parameters throughout large perturbation correction maneuvers. This fitting procedure revealed that the torsional spring rest angle (Fig. 4d) and the damping factor (Fig. 4e) changed significantly during the correction sequence. The torsional stiffness showed only minor changes (Fig. 4f). The same parameter changes were observed across the entire control population (Fig. 4g–i), demonstrating a robust and consistent corrective mechanism captured by the torsional spring model.

**Figure 4.**
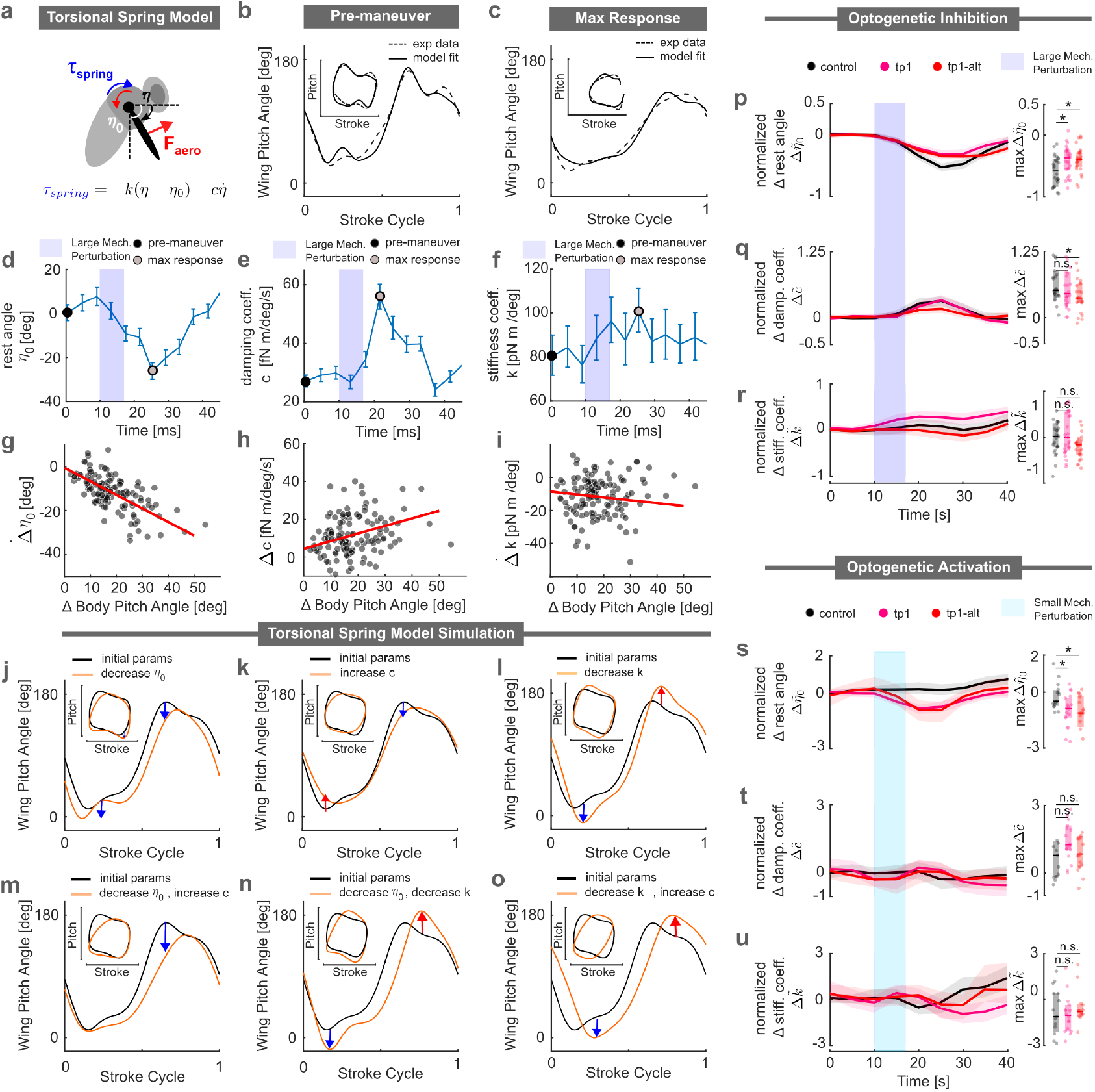
The tp1 muscle modulates the rest angle in a torsional spring model for wing pitch control. (a) Schematic of the torsional spring model. (b) Example from a single fly showing the torsional model fit (solid curve) to pre-maneuver wing kinematics (dashed curve); inset shows the relationship between wing stroke and wing pitch. (c) Example from a single fly showing the torsional model fit (solid curve) wing kinematics associated with maximum change in wing pitch response(dashed curve); inset shows the relationship between wing stroke and wing pitch. (d) Changes in the torsional spring rest angle for the same fly shown in (b) and (c) during large perturbations; the shaded region marks the perturbation period, with black and gray circles indicating the rest angle values corresponding to the conditions in (b) and (c), respectively. (e) Changes in the damping factor for the same fly shown in (b) and (c) during large perturbations. (f) Changes in the stiffness constant for the same fly shown in (b) and (c) during large perturbations. (g–i) Maximum changes in rest angle (g), damping factor (h), and stiffness constant (i) as a function of body deflection amplitude for all control flies (*n* = 164). (j-l) Wing pitch profiles as functions of stroke cycle, comparing reference spring parameters (*k* = 50 pNm/deg, *c* = 10 fNm/deg/s, *η*_0_ = 0 deg) with modified *η*_0_ = 30 deg (j), *c* = 20 fNm/(deg s^−1^) (k), and *k* = 20 pNm/deg (l). The insets illustrate the relationship between wing stroke and wing pitch. (m-o) Combined effects of modifying two spring parameters on wing pitch profiles, with reference parameters in black. Note that the wing pitch profile and inset for (m) closely resemble those for the max response in (c). (p-u) Left: Normalized changes in torsional spring parameters in response to pitch perturbations. Right: Maximum change in normalized torsional spring parameters: - (p-r) Comparison of control and tp1-inhibited flies during large perturbations. -(s-u) Comparison of control and tp1-activated flies during small perturbations. Statistical significance for (p)-(u) is determined via the Kruskal-Wallis test with Dunn’s post hoc multiple comparisons. (***, p<.001; **, p<.01; *, p<.05). Sample sizes for optogenetic inhibition with large perturbation: control *n*=15, tp1 *n*=26, tp1-alt *n*=19. Sample sizes for optogenetic activation with small perturbation: control, *n*=16, tp1 *n*=17, tp1-alt *n*=11.).

To provide intuition for how changes in the various spring parameters influenced the wing pitch profile, we conducted simplified quasi-steady aerodynamic simulations (see Method J) with parameters that qualitatively replicated real wing pitch kinematics, (*η*_0_, *c, k*) = (50, 10, 0). In addition to examining the wing pitch profile over the course of the stroke cycle, we also present an inset showing wing pitch versus wing stroke. This loop features a prominent hump at the top, representing the overshoot in wing pitch angle toward 180° during the transition from the front stroke to the back stroke, as well as a similar overshoot at the lower right toward 0°during the transition from the back stroke to the front stroke. When the torsional spring rest angle (*η*_0_) was reduced to −30° (Fig. 4j), the top overshoot disappeared, and the bottom overshoot deepened. On the other hand, increasing the damping factor (*c*) to 20 fNm/(deg · s^−1^) (Fig. 4k) reduced both overshoots. Finally, decreasing the torsional stiffness (*k*) by 20 pNm/deg (Fig. 4l) amplified both the top and bottom overshoots. These observations showed that reproducing the experimentally observed loop required modifying at least two spring parameters. Among the three possible combinations of two spring parameters shown in Fig. 4m–o, lowering the rest angle (*η*_0_) and increasing the damping factor (*c*) (Fig. 4m) successfully recreated the experimental loop for large perturbations (Fig. 4b). Specifically, these two parameters acted in concert to decrease the top overshoot while maintaining the overshoot at the bottom right. In contrast, the other pairwise combinations (Fig. 4n,o) resulted in more pronounced overshoots or other loop distortions.

Having identified the relevant spring parameter changes, we investigated their effect when the tp1 motoneuron was inhibited during a large perturbation correction. We found that the change in normalized torsional spring rest angle 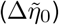 was significantly smaller when the tp1 motoneuron was inhibited compared to controls (red versus black curves in Fig. 4p). We observed no significant differences between the silenced and control groups for the normalized changes in damping factor 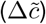 or spring stiffness 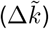 (red versus black curves in Fig. 4q,r). These results indicated that the tp1 muscle was pivotal in modulating the torsional spring rest angle (*η*_0_), one of the two key spring parameters necessary for altering the backstroke wing pitch angle.

Next, we examined the impact of optogenetic activation on torsional spring parameters in the small perturbation regime. We found that activating the tp1 motoneuron significantly altered the normalized torsional spring rest angle 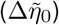 (Fig. 4s). As with the inhibition experiments, however, the normalized changes in damping factor 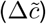 and stiffness constant 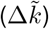 remained unchanged relative to the control group. These findings further supported the idea that the tp1 muscle modulated the rest angle of the wing hinge.

### The tergopleural muscle is modulated by a proportional gain term in a wing pitch PI controller

Finally, to gain an understanding of how the tp1 muscle activity was temporally modulated to produce the observed time course of wing pitch changes during large perturbations, we leveraged a control-theoretic framework established in previous work (Whitehead et al., 2015, 2022). Past studies had shown that pitch stabilization in fruit flies could be effectively modeled using a proportional-integral (PI) controller. In this framework, the controller receives the rate of change of the body pitch angle 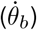 as input, which is sensed by the halteres, the mechanosensory organs that function as vibratory gyroscopes. The controller then output the front wing stroke angle at time *t*, computed as a linear combination of the body pitch angle (Δ*θ*_*b*_) and pitch velocity 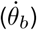 at a previous time *t* − Δ*T* (Whitehead et al., 2022), as illustrated in the orange block of Fig. 5a:

**Figure 5.**
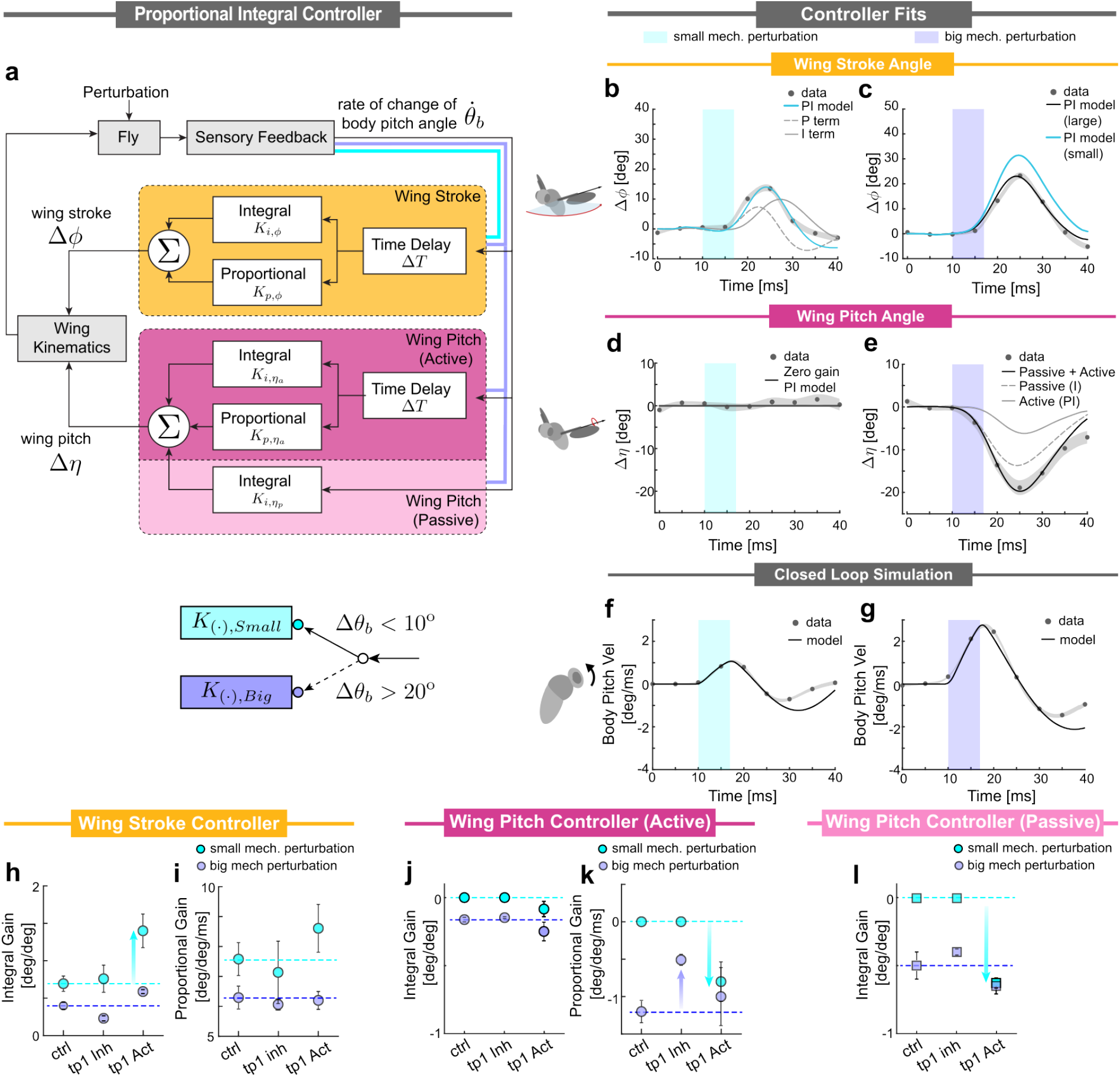
The proportional gain in the pitch stabilization controller modulates tp1 activity. (a) Gain-scheduling Proportional-integral (PI) controller model for rapid flight stabilization in response to the pitch perturbation. The rate of change of body pitch angle 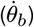 is measured by mechanosensory organs and processed through two parallel controllers: one for wing stroke modulation (orange) and another for wing pitch modulation (pink). In each controller containing the PI motif, the input is subjected to a time delay (Δ*T* = 5 ms) and proportional and integral gains *K*_*p*,·_ and *K*_*i*,·_, respectively. We modeled the observed “preflex” response in wing pitch by incorporating an additional zero-delay integral component (depicted in a lighter shade of pink) into the wing pitch controller. The outputs of these controllers are summed to determine corrective changes in wing kinematics, generating the corrective torque to stabilize the fly. Additionally, the controllers dynamically adjust their gains based on a threshold in the change in body pitch angle, 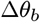 (b,c) Change in wing stroke angle over time for simulated flies (black lines, PI controller) fitted to the experimental data (black circles representing the population average, with the gray shaded region indicating the SEM) for small (b) and large (c) mechanical perturbation. Gray dashed and solid lines in (c) denote the contributions from the P term (proportional), and the I term (integral), respectively. The light blue line in (c) denotes the PI controller model fit with parameters from small perturbation. (d,e) Change in wing pitch angle over time for simulated flies (black lines) fitted to the experimental data (black circles representing the population average, with the gray shaded region indicating the SEM) for small (d) and large (e) perturbation. The solid black line, dashed gray line, and solid gray line in (e) represent the combined passive and active controller model fit, the individual passive model fit, and the active model fit, respectively. (f,g) Change in body pitch velocity over time for simulated flies (black lines) fitted to the experimental data (black circles representing the population average, with the gray shaded region indicating the SEM) for small (f) and large (g) perturbation. (h,i) Summary statistics for wing stroke controller parameters— integral gain (*K*_*i,ϕ*_, h) and proportional gain (*K*_*p,ϕ*_, i) showing the mean*±*SEM for small (light blue) and large (blue) perturbation. (j,k) Summary statistics for wing pitch active controller parameters— integral gain (*K*_*i,η*_) and proportional gain (*K*_*p,η*_) showing the mean*±*SEM. (l) Summary statistics for wing pitch passive controller parameter: integral gain 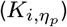 showing the mean*±*SEM. Sample sizes for small perturbation: control (control) *n*=28, optogenetic activation (tp1 act) *n*=37, optogenetic inhibition (tp1 inh) *n*=27. Sample sizes for large perturbation: control *n*=61, tp1 act *n*=43, tp1 inh *n*=53.

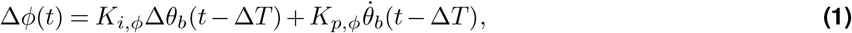

where *K*_*p,ϕ*_, *K*_*i,ϕ*_, and Δ*T* represents the proportional gain, integral gain, and time delay/reflex latency, respectively. The proportional and integral gains determine the relative contributions of angular velocity and displacement to the corrective response, while the time delay corresponds to neural processing and actuation delay in the reflex pathway. In Eq. (1), the body pitch angular displacement (Δ*θ*_*b*_) is defined relative to the fly’s pre-perturbation attitude and can be obtained by integrating the angular velocity signal 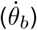. This controller model could also be formulated as a proportional–derivative (PD) controller, with angular displacement as the input. However, previous studies have established that halteres are primarily sensitive to angular velocity rather than angular position, and therefore we adopted the PI convention established in the literature (Whitehead et al., 2022, 2015; Beatus et al., 2015). That said, there is no direct evidence that the angular pitch velocity signal from the haltere is integrated to produce a neural representation of angular displacement; in fact, the fly’s pitch orientation may instead be measured via other sense organs such as the ocelli (Parsons et al., 2010; Krapp, 2009), vision (Kim et al., 2017; Suver et al., 2016; Sugiura and Dickinson, 2009; Erginkaya et al., 2025; Theobald et al., 2010), or the antennae (or legs) (Budick et al., 2007; Mamiya and Dickinson, 2015; Fuller et al., 2014; Mamiya et al., 2011; Mills et al., 2025).

We found that the PI model in Eq. (1) effectively captured the wing stroke response for the small perturbation stabilization reflex (Fig. 5b). The same controller gains, however, failed to accurately reproduce experimental wing stroke data for large perturbations (light blue curve, Fig. 5c). This discrepancy suggests that a fixed-gain controller is insufficient and that a gain-scheduling approach, where the controller gains depend on the perturbation regime, may better capture the observed dynamics (Fig. 5a bottom, Supplementary Text). Once the controller gain is adjusted to account for this difference in regime, we find that the wing stroke response is well captured by the controller (black line, Fig. 5c).

To account for the changes in the wing pitch angles we introduce an additional control block (pink block in Fig. 5a)—which regulates wing pitch based on the rate of change of the body pitch angle. For small pitch perturbations (light blue pathway in Fig. 5a), this block is inactive (Fig. 5d). For the large perturbation regime, this controller features a PI architecture with a time delay, consistent with that for stroke control. Interestingly, careful inspection of the wing pitch modulation shows an instantaneous response component as indicated by a decrease in wing pitch that is nearly coincident with the onset of the mechanical perturbation. This observation suggests the possibility of a passive, mechanically mediated adjustment to the wing pitch—often referred to as a ‘preflex’ (Jindrich and Full, 2002; Proctor and Holmes, 2010)—that occurs without the latency of neural processing. To capture this effect, we incorporate an additional zero-delay integral term (light pink block in Fig. 5a). The combined controller can be expressed as:

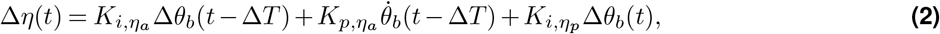

Where 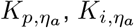, and Δ*T* represent the proportional gain, integral gain, and time delay of the active wing pitch controller, respectively. The last term in Eq. (2) with integral gain, 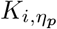 represents the zero-delay (putatively passive) component. We find that this wing pitch controller, which combines active (gray solid line) and passive (gray dashed line) blocks, accurately captures the wing pitch response in the large perturbation regime (black line in Fig. 5e).

To further validate the robustness of this combined controller framework, we performed closed-loop simulations of a fly experiencing a pitch perturbation and implementing a correction maneuver guided by these controllers. Here, we use a linearized form of the body pitch dynamics along with quasi-steady aerodynamic calculations to estimate the torques generated by the wing strokes. The simulation results for the body pitch velocities as a function of time for the two perturbation regimes are shown in Fig. 5f,g. We find quantitative agreement between the experimental measurements (data points) and simulation results (black lines), particularly in the early part of the correction maneuver, further confirming the accuracy of these controllers.

This combined controller model provides a useful framework for understanding how the underlying reflex circuit modulates the tp1 muscles, which are critical for implementing the large perturbation response. Specifically, we next investigated how the wing stroke, the active wing pitch, and the low-latency (passive) wing pitch controller fits are influenced by optogenetic manipulations of the tp1 motoneuron (Fig. 5h-l). For each condition, we fit the wing stroke responses to the controller model and extract the corresponding controller gains. Consistently, across all experimental conditions—including genetic control, and optogenetic activation and inhibition—we observed a systematic decrease in the fitted controller gains, *K*_*i,ϕ*_ and *K*_*p,ϕ*_ as we transition from the small perturbation regime (light blue data) to the large perturbation regime (purple data). This trend indicates that the controller has two distinct operating modes. We first focus on the orange block of Fig. 5a, corresponding to the wing stroke controller. We find that optogenetic inhibition of the tp1 motoneuron only had minor effects on the integral gain in both perturbation regimes. In contrast, tp1 activation had a large effect on the integral gain in the small perturbation regime (light blue arrow in Fig. 5h). We observed only minor effects due to inhibition and activation on the proportional gain (Fig. 5i). These data indicate that activation of the tp1 muscle boosts the effect of the integral gain portion of the controller for wing stroke modulation.

Next, we focus on the pink block in Fig. 5a, corresponding to the wing pitch controller. We observed that optogenetic inhibition and activation of the tp1 motoneuron had only minor effects on the active block controller’s integral gain across both perturbation regimes (Fig. 5j). In contrast, while tp1 inhibition did not affect the active block controller’s proportional gain in the small perturbation regime, it significantly reduced this gain in the large perturbation regime (light purple arrow in Fig. 5k). Conversely, tp1 activation in the small perturbation regime markedly increased the magnitude of the active block proportional gain (light blue arrow in Fig. 5k), while having only a marginal effect in the large perturbation regime. Since both silencing and activation of the tp1 motoneuron alters the active block proportional gain, these results suggest that the output of the proportional gain may be modulating the tp1 motoneuron, which in turn drive changes to the wing pitch.

Finally, we note that tp1 inhibition had minimal impact on the passive block integral gain across both perturbation regimes. Interestingly, tp1 activation led to an increase in passive integral gain during the small perturbation regime. We speculate that this shift may arise from changes in thoracic or wing mechanics induced by the unnatural activation of the tp1 motoneuron in the small perturbation regime.

## Discussion

In this study, we demonstrated that Drosophila exhibit strength-dependent, compensatory strategies to recover from pitch disturbances. Through optogenetic manipulation of its motorneuron, we found that the tp1 muscle is recruited in response to large, but not small, perturbations, apparently playing a role in modulating wing pitch angle. Motivated by this insight, we extended an existing control-theoretic framework (Whitehead et al., 2015, 2022; Beatus et al., 2015; Ristroph et al., 2010) to more accurately capture the fly’s corrective responses and found that the wing pitch dynamics are modulated by the proportional component of a PI controller. Using a simplified model of wing motion in which the base of the wing is represented as a torsion spring that deforms under the instantaneous aerodynamic torque (Beatus and Cohen, 2015; Bergou et al., 2007), we found that tp1 activity was correlated with adjustments to the rest angle for wing pitch. Together, these results suggest that neuromuscular recruitment during aerial correction maps onto distinct components of canonical feedback control structures.

Interestingly, we also found that the correction strategies observed in our experiments could be closely reproduced by simulating wing kinematic adjustments under a power minimization principle. Although speculative, this finding suggests that energetic considerations may act as an additional constraint in shaping the neuromuscular control of flight. Our data do not demonstrate that this reflex pathway evolved explicitly to minimize energetic cost, but the correspondence between observed responses and power-minimizing predictions is consistent with a broader pattern in locomotor systems, where control strategies often converge on energetically favorable solutions (Margaria, 1938; Griffin et al., 2004; Brown et al., 2021; Donelan et al., 2001; Ralston, 1958; Hoyt and Taylor, 1981). Given that flapping flight is among the most energetically costly modes of locomotion, it remains plausible that energy efficiency places meaningful constraints on how pitch correction is implemented.

Our torsional spring model provides a useful and analytically tractable framework for understanding how tp1 activity might influence wing pitch. However, it necessarily abstracts away the substantial mechanical complexity of the wing hinge, which has been the focus of decades of biomechanical study. Such studies have shown that the primary rotational joint of the wing is formed by the second axillary sclerite (ax2) and its ventral articulation with the pleural wing process (pwp), which serves as the fulcrum for wing motion (Boettiger and Furshpan, 1952; Pringle, 1957; Miyan and Ewing, 1985; Walker et al., 2014; Deora et al., 2017; Melis et al., 2024). Distally, ax2 is firmly connected to the wing’s main structural vein (the radius), making its orientation critical for determining the angle of attack. Boettiger and Furshpan (1952) first proposed that consistent changes in angle of attack throughout the stroke result from a phase difference between the anterior notal wing process, which actuates ax2 anterior to its articulation with the pwp, and the scutellar lever arm, which actuates ax2 posterior to this articulation. In this view, the overall pattern of wing rotation is mechanically embedded in the hinge exoskeleton by the indirect power muscles, with fine-scale adjustments mediated by the direct steering muscles (Lindsay et al., 2017; Melis et al., 2024). Although a resilin-rich flexion joint at the wing base has been identified that allows for passive modulation of stroke amplitude under inertial load at stroke reversal (Lerch et al., 2020; Melis et al., 2024), no evidence has been found for any structure that on its own could function as a torsional spring specifically regulating wing pitch. In this light, the torsional spring framework should be regarded as capturing a relatively simple emergent behavior of a very complex joint. The potential role of tp1 within this framework remains unresolved. In Drosophila, genetic silencing experiments have suggested a contribution to male courtship song rather than to flight directly(O’Sullivan et al., 2018). Anatomically, tp1 does not insert on any of the four axillary sclerites of the wing but instead makes broad contact with the mesoscutum near the convergence of the transverse and notopleural ridges (Wisser and Nachtigall, 1984))From its morphology, it seems unlikely that tp1 directly sets the wing-pitch rest angle. Nonetheless, given the overall complexity of the wing hinge, it is possible that tp1 contributes indirectly (Dickinson and Tu, 1997)—for example, by subtly deforming the mesoscutum and thereby modifying the tension among all the wing hinge sclerites and the muscles attached to them. Such alterations within the hinge might modify wing pitch kinematics in a manner consistent with the rest-angle adjustments posited by the simplified model, thus shaping higher-order control dynamics.

Another important open question arising from our study concerns the functional role of the tp1 muscle during unperturbed, free flight. Specifically, does optogenetic manipulation of tp1 motoneurons alter wing pitch modulation in the absence of external perturbations, or does tp1 primarily act to amplify the effects of other muscles? To investigate this question, we conducted free-flight optogenetic experiments (Supp. Fig. S9). Inhibition of tp1 produced no statistically significant changes in body or wing kinematics, suggesting that tp1 is not required for baseline flight control. Conversely, optogenetic activation led to a modest decrease in vertical body velocity and subtle front stroke changes, but had no significant effect on wing pitch angle. These results imply that tp1 alone may be insufficient to modulate wing pitch dynamics and may instead serve as a modulatory or amplifying component within a broader motor program. Previous morphological and functional studies (Miyan and Ewing, 1985; Dickinson and Tu, 1997; Lindsay et al., 2017; Melis et al., 2024) highlight the iv (also known as hg) muscles—which attach to the fourth axillary sclerite—as strong candidates for directly driving wing pitch rotations. Future investigations that simultaneously manipulate tp1 and iv muscles may clarify whether these muscles act cooperatively to control wing kinematics, particularly during high-demand corrective maneuvers.

Our control-theoretic analysis revealed a reduction in stroke controller gains during large perturbations (Fig.5h–i). While this reduction in gain could reflect changes in neural circuitry, it may also arise from nonlinearities in muscle power output. Prior work has shown that muscles in other animals often operate near the steepest portion of the power–phase activation curve, where small shifts in activation timing yield large changes in mechanical output (Sponberg and Daniel, 2012; Tu and Daniel, 2004; Josephson, 1985; Ahn and Full, 2002; Marsh and Olson, 1994; Ettema, 1996; D’Aout et al., 2001; Girgenrath and Marsh, 1999). This steep regime enables precise control via phase modulation, particularly during unsteady maneuvers. Large perturbations, however, may shift muscle activation away from this optimal phase range into a saturated regime, where output becomes less sensitive to timing. This shift flattens the phase–power relationship, effectively reducing the system’s responsiveness and lowering controller gains. Thus, phase-dependent muscle nonlinearity provides a plausible mechanistic basis for the observed drop in control gain during extreme maneuvers.

While muscle-level nonlinearities may explain the reduction in stroke controller gains, the onset of the wing pitch response likely originates from distinct neural mechanisms. One plausible mechanism involves coincidence detection among primary sensory afferents—especially those from the halteres, which serve as gyroscopic sensors of body rotation (Pringle, 1948; Yarger and Fox, 2018; Mohren et al., 2019). These afferents may require temporally aligned activation across multiple sensilla to drive downstream interneurons, resulting in a dependence on perturbation strength. For example, stronger perturbations could either recruit more haltere afferents or increase the synchrony of their spiking activity, either of which could increase the likelihood of reaching the excitation threshold for tp1 motoneuron activation. Such a mechanism would function as a neural switch, to activate pitch corrections when perturbations are large. This mechanism could explain the absence of tp1 engagement during steady or mildly perturbed flight and its selective activation during larger maneuvers. Future electrophysiological studies will be essential to determine whether such nonlinear integration underlies the sensorimotor transformation driving wing pitch control. Recent work has begun to explore the underlying neural pathways involved in flight control and haltere feedback, laying the groundwork for probing these mechanisms in more detail (Dickerson et al., 2019; Verbe et al., 2024; Dhawan et al., 2025)

Collectively, this study revealed a functional role for the tp1 indirect wing-steering muscle during flight stabilization that is distinct from its role during courtship (von Philipsborn et al., 2011; Ehrhardt et al., 2023). In particular, our conjecture that the tp1 muscle enhances the response of other steering muscles is quite intuitive and consistent with our everyday experience. As anyone who has ever tripped while walking will recall, many muscles beyond those directly controlling the legs are activated during the recovery. Our novel findings shed light on how *Drosophila* are able to leverage the beautiful interplay between such indirect and direct steering muscles to produce their emergent elegant corrective responses to a midair trip.

## Methods

### A. Fly stocks and fly handling

Flies used for opto-mechanical experiments were reared in the dark at room temperature on 0.4mM retinal food (Media Facility, HHMI Janelia Research Campus). Wild-type files used for mechanical perturbation were raised at room temperature on a standard fly medium made from yeast, agar, and sucrose with a 12-hour light/12-hour dark cycle. Female flies, 4 to 6 days after eclosion, were used for all flight experiments. A full list of *Drosophila melanogaster* stocks used in this paper is given in Table 1.

**Table 1.**
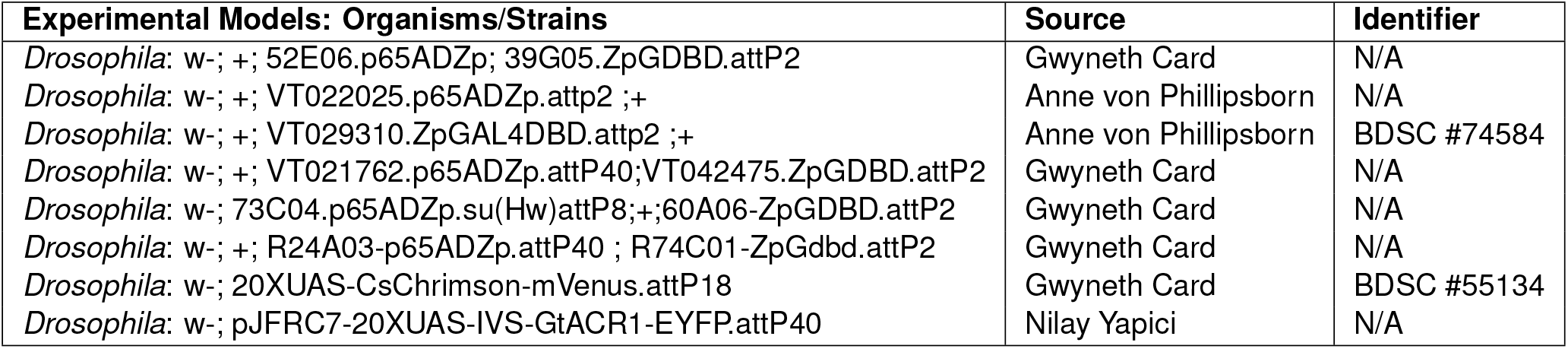
Fly stocks.

### B. Immunohistochemistry

To visualize the nervous system, we dissected *Drosophila* CNS in PBS using fine forceps and placed them in 4% paraformaldehyde solution (in PBS) for 15 min at room temperature. Subsequently, we washed the CNS in PBST three times, 15 minutes each, before incubating the CNS in normal goat serum (NGS) for 20 minutes. We stained the tissue overnight at 4°C with 1:10 mouse anti-nc82 (DSHB) and 1:1000 rabbit anti-GFP (Invitrogen no. A6455) in PBST. Then, we washed the CNS in PBST and applied a secondary antibody stain consisting of 1:250 goat anti-mouse AlexaFluor 633 (Thermo Fisher no. A21052) and 1:250 goat anti-rabbit AlexaFluor 488 (Thermo Fisher no. A11034) in PBST overnight at 4°C. We then removed the CNS from the solution and placed them in PBS for 30 min on an oscillator at room temperature. Afterward, we mounted the CNS on a glass microscope slide in Vectashield (Vector Labs H-1000-10), covered with one #1.5, 22 × 22 mm coverslip, and sealed with nail polish. We imaged the CNS at 20x magnification on a Zeiss LSM 880 confocal microscope at 1 *µ*M depth resolution. Final images are presented at maximum z-projections over relevant depths.

To visualize the muscles and their innervations, flies were rinsed in 95% ethanol before they were then frozen for 3-4 min at −20° C in a mold containing Tissue-Tek O.C.T. compound (VWR Chemicals). The frozen thoraxes were then hemisected along the midline and transferred into 4% paraformaldehyde in PBS for 30 mins. Samples were incubated with 1:10 mouse anti-nc82 (DSHB), 1:1000 rabbit anti-GFP (Invitrogen no. A6455), and Alexa 568 phalloidin (Thermo Fisher Scientific, 1:500) at 4°C for 3-4 days. Then, we washed the CNS in PBST and applied a secondary antibody stain consisting of 1:250 goat anti-mouse AlexaFluor 633 (Thermo Fisher no. A21052) and 1:250 goat anti-rabbit AlexaFluor 488 (Thermo Fisher no. A11034) in PBST overnight at 4°C for 5-6 days. We mounted the hemisected thoraxes on a glass microscope slide in Vectashield (Vector Labs H-1000-10), covered with one #0, 22 × 22 mm coverslip, and sealed with nail polish. We imaged the CNS at 20x magnification on a Zeiss LSM 880 confocal microscope at 1 *µ*M depth resolution. Final images are presented at maximum z-projections over relevant depths.

### C. Fly preparation

Individual flies were anesthetized on a custom-made cold plate, regulated by Peltier devices, with temperatures ranging from 0°C to 4°C. Subsequently, a ferromagnetic pin, measuring between 1.4 and 2 mm in length and 0.15 mm in diameter, was affixed to the flies’ notum along their sagittal plane using UV glue. The added mass of the pins is comparable to the natural mass variation of the flies and negligibly impacts the off-diagonal components of the fly’s inertia tensor. Experiments conducted with unpinned flies in free flight demonstrated that the addition of a pin did not qualitatively affect the flies’ flight kinematics. For detailed calculations, refer to (Whitehead et al., 2015; Beatus et al., 2015).

### D. High-speed videography

We conducted experiments utilizing 10 to 25 flies per genotype, following the aforementioned fly preparation procedure. These flies were introduced into a custom-built cubic flight chamber measuring 13 cm on each side. Positioned at the chamber’s center were three orthogonal high-speed cameras (Phantom V7.1), capturing footage at 8000 frames per second with a resolution of 704 × 704 pixels. Illumination for each camera was provided by near-infrared light-emitting diodes (LEDs) (850 ± 30 nm, Osram Platinum Dragon). Flies’ entry into the filming zone of the high-speed cameras was detected via an optical trigger system (Ristroph et al., 2009). This system utilized split and expanded beams from a 5-mW, 633-nm HeNe laser (Thorlabs, HRR050), passing through a neutral density filter (Thorlabs, NE20A) with an optical density of 2.0 before reaching two photodiodes (Thorlabs, FDS100). Prior to each experimental session, camera calibration was performed using the easyWand protocol detailed in (Theriault et al., 2014).

### E. Opto-mechanical perturbation experiments

For each optomechanical perturbation experiment, a cohort of 10 to 25 flies, each affixed with a pin to their notum, was released into the flight chamber as previously described, remaining therein for approximately 12 hours. To apply midair optomechanical perturbations, we used the optical trigger circuit outlined earlier, which drove a 45-ms duration voltage pulse to an LED driver (Thorlabs, LEDD1B). This LED driver then supplied a 1-A current to either a 625 nm red LED (Thorlabs, M625L4) or a 565 nm green LED (Thorlabs, M565L3) for optogenetic excitation (CsChrimson) or inhibition (GtACR1) experiments, respectively. Both red and green LED sources were equipped with a collimating attachment (Thorlabs, COP2-A) to produce a 50-mm-diameter beam profile. The resulting intensities were 731 and 316 *µ*W/mm^2^ for the red and green LEDs, respectively. The cross-sectional area of this beam was sufficiently large to ensure that the light source would unavoidably intersect a fly anywhere within the filming volume of all three cameras. Additionally, the collimation ensured uniform stimulus intensity irrespective of the fly’s position within the filming volume. To prevent external light contamination during these experiments, the entire flight apparatus was enclosed by blackout curtains. Given that flies are unlikely to initiate flight bouts in total darkness, a dim, blue fluorescent light bulb illuminated the arena during experimental trials. This protocol was also used for optogenetics experiments.

Following a 10ms delay, the trigger initiated a 7ms magnetic field pulse by delivering a rapid current pulse to a pair of Helmholtz coils positioned on the top and bottom faces of the flight chamber. The placement of these coils generated a roughly uniform vertical magnetic field in the center of the filming volume, activated upon a fly’s entry into this region of the flight chamber. Typical magnetic field strengths were on the order of 10^−2^ T. The magnetic field from the coils acts on the magnetic moment of the ferromagnetic pin glued to the fly, consequently inducing a moment about the fly’s pitch axis. To generate pitch perturbation of different magnitudes, the voltage source supplied to the Helmholtz coils was varied from 15V to 28V. Further details of this procedure are described in (Whitehead et al., 2015, 2022; Ristroph et al., 2010; Beatus et al., 2015). Simultaneously; the high-speed cameras recorded flight activity at 8000 frames per second, encompassing periods before, during, and after the optogenetic stimulation and application of the magnetic field.

### F. Flight data selection and kinematic extraction

From the data collected in both optogenetic and optomechanical perturbation experiments, we initially filtered the fly movies to ensure that each contained a fly within the field of view of all three cameras within the time range of *t* ∈ [−10, 40]ms, facilitating further analysis. In addition, we focused our attention only on videos where there was no evidence of a fly performing volitional maneuvers before the onset of optogenetic stimulation and mechanical perturbation. Given that our optogenetic activation/inhibition, lasting 45 ms, commences at *t* = 0ms and mechanical perturbation begins at *t* = 10ms, we imposed additional criteria aimed at videos where the mechanical perturbation primarily influences the body’s pitch axis. This criterion was applied to isolate corrective maneuvers related to a single rotational degree of freedom.

To extract kinematic data from the three high-speed camera views, we used a custom 3D hull reconstruction algorithm detailed in (Ristroph et al., 2009). This algorithm allows us to extract 12 kinematics variables describing the orientation of a fly - the 3D position of the fly center of mass and three sets of Euler angles describing the fly body, left and right-wing-at each time point. In the event of encountering an occlusion that prevents a direct extraction of a particular kinematics variable, we estimate the missing data values by using a cubic spline interpolation. As the wing response to our perturbation is symmetric, we select one of the wings with the best-estimated wing span and wing chord vector from the wing visual hull for further analysis to constrain wing angle estimation errors due to occlusion. In our analyses, similar to (Whitehead et al., 2022), we filtered the raw body kinematics with a 100Hz low-pass filter. We smoothed the raw wing kinematics using the Savitzky-Golay method. In particular, we used a 7th-order polynomial with a window size of 21 frames (2.625 ms) to smooth the wing stroke angle; for the wing deviation and rotation angles, we used a 5th-order polynomial and a window size of 11 frames (1.375 ms).

To examine the effect of optogenetic manipulation of tps motoneuron on body kinematics, as showcased in Fig. 1 (g,h) and 3 (a,d,g,j), we aligned the body pitch angle time series to a standardized time interval [0,40ms] by uniformly resampling the time series via a spline fit (MATLAB’s smoothing spline function spaps.m). The resampled time series were subtracted from their baseline, defined as the average body pitch angle from 0ms to 10ms. Subsequently, the body pitch kinematics were normalized by the maximum deflection amplitude before the population responses were quantified via the average of the time series.

To average wingbeat kinematics across flies—as in Fig. 2 (b,d,f,h)—we isolated wingbeats from time series data using stroke angle maxima as reference points. Segmented wingbeat kinematics were then aligned to a standardized wingbeat cycle time by uniformly resampling Euler angle values via a spline fit (MATLAB’s smoothing spline function spaps.m). For extracting the maximal wing response to pitch perturbation, depicted in Fig. 2 (i,j), we identified the stroke angle minimum and wing pitch angle maximum within each wingbeat. The maximal wing responses were then located from the time series of wing features subtracted from the baseline estimate, defined as the averaged wing feature over the pre-perturbation period (0ms to 10ms).

To evaluate the population-wide wing response to pitch perturbation, normalized by the perturbation strength, as depicted in Fig. 2 (i,j) and Fig. 3 (b,c,e,f,h,i,k,l), we employed the same approach as outlined above, identifying the front stroke angle and backstroke wing pitch angle response from the baseline. Subsequently, the wing responses were normalized by the maximum deflection amplitude before employing a windowed average estimate with a window size of 2.5ms.

### G. Backstroke wing pitch response curve fitting

To model our backstroke wing pitch response, Δ*η* to the strength of deflection amplitude Δ*θ*_*B*_, we considered both a linear fit, Δ*η*(Δ*θ*_*B*_ | *β*_0_, *β*_1_) = *β*_1_Δ*θ*_*B*_ + *β*_0_ and a sigmoidal fit, Δ*η*(Δ*θ*_*B*_ | *η*_*max*_, *α*, Δ*θ*_0_) = Δ*η*_*max*_(1 +exp(*α*(Δ*θ*_*B*_ − Δ*θ*_0_)))^−1^. We compared the two models based on Akaike Information Criterion (AIC) (Akaike, 1987; Anderson et al., 1998; Bozdogan, 1987), AIC = log MSE + 2*m* + 2*m*(*m* + 1)*/*(*n* − *m* − 1), where MSE is the mean squared error from our least square fitting, *m* is the number of parameters and *n* is the number of data points. AIC provides a mean for model selection based on the relative goodness of fit of different models while accounting for the number of parameters. We selected the sigmoidal model based on its lower AIC score (Sigmoid model AIC : 686.427 < Linear model AIC : 694.114).

### H. Quasi-steady aerodynamic calculation

To estimate the aerodynamic forces and torques generated by the wing kinematics observed in our experiment, we utilize a quasi-steady aerodynamic model developed in previous studies validated with scaled mechanical models(Sane and Dickinson, 2002; Whitney and Wood, 2010; Dickinson et al., 1999). In particular, we calculated the total instantaneous quasi-steady aerodynamic force on a wing :

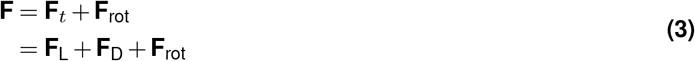

where **F**_L_ and **F**_D_ are the lift and drag components of the translational force **F**_*t*_, and **F**_rot_ is the rotational force. Similar to (Fry et al., 2005; Whitehead et al., 2022), We did not include the contribution from the added-mass effect owing to the large error in estimating the second derivative of the angle of attack from experimental data. Their full expressions are :

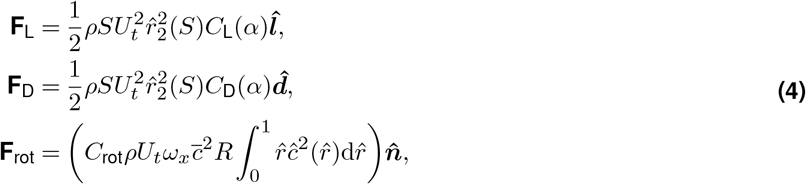

where *ρ* is the air density; *S* is the wing area; *U*_*t*_ is the velocity of the wing tip; 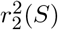 is the non-dimensionalized second moment of wing area; *α* is the wing angle of attack; *C*_L_ and *C*_D_ are the lift and drag coefficients; *C*_rot_ the is rotational force coefficient; *ω*_*x*_ is the angular velocity of the wing about its spanwise axis; *c* is the mean wing chord length; *R* is the wing span length; 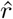 and *ĉ* the non-dimensionalized span and chord; and 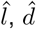, and 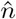 are unit vectors in the directions of the lift, drag, and wing surface normal. The *C*_L_ and *C*_D_ coefficients are given by :

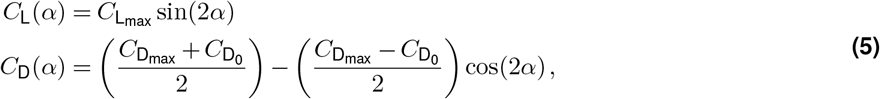

where 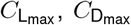, and 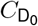 are dimensionless parameters best fit for *Drosophila* flight. We summarized all the values used in our computation in Supp. Table. 1.

To calculate the aerodynamic torque, **T**_aero_ produced by a flapping wing, we approximate the wing center of pressure to be 70% along the wing span (Birch and Dickinson, 2001):

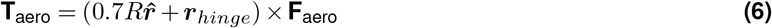

where R is the wing span length, 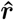 is the wing span unit vector, **F**_aero_ is the total aerodynamic force produced by the wing, and ***r***_*hinge*_ is the vector from the fly center of mass to the wing hinge. Similar to (Whitehead et al., 2015, 2022), as we are primarily concerned about the pitching torque, we express the wing hinge vector to be purely along the fly’s long body axis, 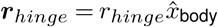.

To determine the power required to generate a particular set of desired wing kinematics, we estimate the power necessary to overcome both aerodynamic **T**_aero,w_ and inertial torques **T**_inertial,w_ in the wing frame of reference:

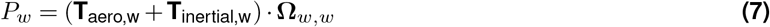

where **Ω**_*w,w*_ is the angular velocity vector of the wing in the wing frame of reference. Similar to (Berman and Wang, 2007), we assume that the cost for negative power is negligible and that the effect of elastic storage is minimal.

### I. Modeling the pitch correction response

To model the selection strategy for pitch correction described in the main text, we utilized representative pre-maneuver wingbeat kinematics and corrective wingbeat kinematics to sample possible wing modulations during a corrective response to pitch perturbations. The transition between pre-maneuver and corrective wing responses was modeled using linear interpolation:

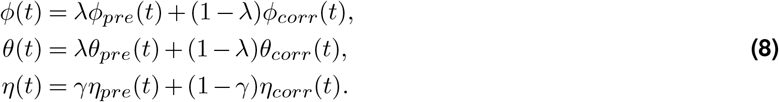

where *λ* controls the interpolation for wing stroke *ϕ* and wing deviation *θ* as they are strongly coupled, and *γ* controls the interpolation for wing pitch *η*. As the deviation angle response is always correlated to the wing stroke response, we use the same *λ* as the wing stroke. Using this interpolation approach, we densely sampled pitch torques *τ* and lift forces *F*_*L*_ across the range of wing kinematics observed in our experiments. For simplicity, we assumed that a fly could correct a pitch perturbation and return to its initial orientation by generating a corrective pitch torque, 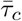, over two wingbeats. To generate a specific corrective torque, 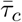, we identified an isoline that contains a series of wing modulations, 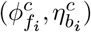, that achieved 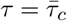. We then traversed along the isoline by progressively adjusting the modulations, changing both front stroke and backstroke wing pitch, until we identified a combination, 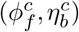, that satisfies the lift maximization or power minimization constraints. This procedure was repeated over various pitch torques to explore the corresponding wing modulations under the two constraints.

### J. Torsional spring model fitting and simulation

We followed the approach outlined in (Beatus and Cohen, 2015) to estimate the torsional spring parameters best fitting the experimentally observed wing pitch angle. Briefly, for each wingbeat, the body angular orientation (*ϕ*_*B*_, *θ*_*B*_, *ψ*_*B*_), body center of mass velocity (*v*_*x*_, *v*_*y*_, *v*_*z*_), wing stroke *ϕ*, wing deviation *θ* kinematics, and an initial condition for the wing pitch angle 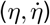 were combined with a set of torsional spring parameters (*η*_0_, *k, c*) to solve the wing pitch angle’s equation of motion,

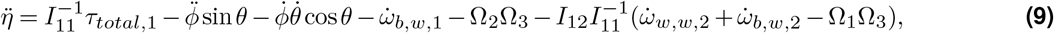

where *I*_1,1_ and *I*_1,2_ are entries of the wing moment of inertia tensor, *τ*_*total*,1_ is the first component of the sum of aerodynamic and spring torques, Ω_1_, Ω_2_ and Ω_3_ are the wing angular velocity vector component in the wing frame of reference, 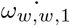 is the first component of the body angular acceleration in the wing reference frame, 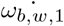 is the first component of the wing angular acceleration in the wing reference frame. The equation was solved using MATLAB’s ODE solver ode45. To identify the optimal spring parameters, we performed a grid search over the torsional spring parameter space, minimizing the root mean squared error (RMSE) between the calculated and experimentally measured wing pitch angles.

To examine the influence of spring parameters on wing pitch kinematics (Fig. 4j-o), we assumed a body angular orientation of (0, *π/*4, 0), zero body center of mass velocity, and simplified wing kinematics: *ϕ*(*t*) = *A* sin(*ωt*) −*π/*2 with *A* = 4*π/*9 for the wing stroke and *θ*(*t*) = 0 for the wing deviation. Using these assumptions, we varied the spring parameters while solving Eq. 9 to compute the resulting wing pitch kinematics.

To examine the influence of tp1 motoneuron on the spring parameters, —as in Fig. 4 (p-u), we extracted the spring parameters for each wing beat. The spring parameters response was then subtracted from the baseline estimate, defined as the averaged spring parameters over the pre-perturbation period (0ms to 10ms). The spring parameters were then normalized by the maximum deflection amplitude before employing a windowed average estimate with a window size of 2.5ms.

### K. Control theory model fitting

For the control theory model fitting, we combined the data across all genetic lines for each condition (control, optogenetic activation, and inhibition), as the inter-line variance was negligible. We estimated controller transfer functions and hence the controller paramters using MATLAB’s tfest command, with body pitch velocity as the input and wing stroke and pitch profiles as the outputs. For tfestOptions, we applied the Levenberg-Marquardt least-squares method with ‘InitializeMethod’ set to ‘all’, ‘EnforceStability’ set to ‘true’, and ‘InitialCondition’ set to ‘Estimate’. For the assessment of fitting quality, we used:

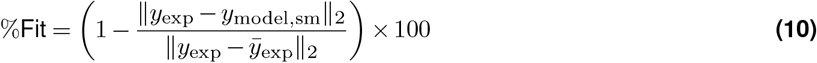

where *y*_exp_ is the experimental data (wing stroke/pitch profiles), *y*_model,sm_ is the respective model fit smoothened with a moving average filter of window length of 2.6 ms, 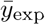 is the mean of the experimental data,*y*_exp_, and ||·||_2_ is the *L*_2_ norm.

For modeling wing kinematics and fly mechanics, we adopted a similar approach to our previous study (Whitehead et al., 2022). We approximate the output of the wing kinematics block, the pitch torque (*T*_pitch_), as a linear combination of the differential wing stroke (Δ*ϕ*) and wing pitch angle (Δ*η*) as follows:

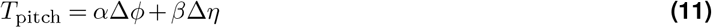

where the coefficients are *α* = 0.006629 and *β* = −0.002207. The negative sign of *β* aligns with the observed decrease in differential wing pitch. For the linearized flight dynamics, we assume:

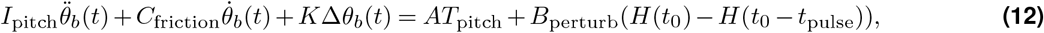

where *I*_pitch_ is the pitch moment of inertia, *C*_friction_ is the coefficient of pitch rotational drag (Supplementary Table 3). 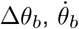, and 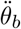 are the body pitch angle, velocity, and acceleration, respectively. *H* represents the Heaviside step function, which is nonzero only during the duration of the applied perturbation, *t* ∈ [*t*_0_, *t*_0_ + *T*_pulse_] with *t*_0_ = 10 ms and *T*_pulse_ = 7 ms. The first fitting parameter *K* is analogous to a spring constant that captures the relationship between body pitch angle and torque that can arise, for example, due to an offset between the center of lift and center of mass (Ristroph et al., 2010; Beatus et al., 2015). The second fitting parameter, *A*, captures the scaling that relates changes in wing kinematic output to changes in pitch torque depending on the perturbation strength and the fly lines used.

This linearized model leads to the following transfer functions for the plant (*P* (*s*)) and composite proportional-integral controller (*C*(*s*)) as follows:

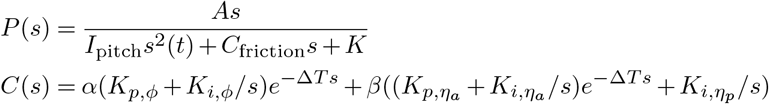

Here, *P* (*s*) is the transfer function from *T*_pitch_ to the angular velocity. where *s* is the Laplace transform complex frequency and *e*^−Δ*T s*^ represents the controller time delay (5 ms for control, tp1; 4 ms for tp2 and tpn), *K*_*p*,·_ and *K*_*i*,·_ are the proportional and integral gain parameters of the respective controller. Strictly speaking, the integral feedback need not be computed by integrating angular velocity, but may instead arise from a direct measurement of pitch angle from the compound eyes (Kim et al., 2017; Suver et al., 2016; Sugiura and Dickinson, 2009; Erginkaya et al., 2025; Theobald et al., 2010), antennae (Budick et al., 2007; Mamiya and Dickinson, 2015; Fuller et al., 2014; Mamiya et al., 2011; Mills et al., 2025), or ocelli (Parsons et al., 2010; Krapp, 2009). Using MATLAB’s feedback command, we generated the closed-loop transfer function *G*(*s*) = *P* (*s*)*/*(1 + *P* (*s*)*C*(*s*)). where *G*(*s*) maps pitch perturbation, *B* to pitch angular velocities. The mechanical perturbation, *B* is modeled as a resistance-inductance (RL) circuit current profile with a time constant of 0.5 ms, where the perturbation is applied starting at 10 ms and terminated at 17 ms.

## Acknowledgment

We thank Anne von Philipsborn for providing us with UAS-Chrimson; VT22025-p65.ADZ VT29310-GAL4.DBD for the tp1-SG experiments. We are grateful to Brad Dickerson’s lab for sharing the muscle hemisection protocol. We also would like to thank Marie Suver’s lab for sharing the CNS dissection and imaging protocol. H.K. Teoh was supported in part by a Mong Junior Cornell Neurotech Fellowship. H.K. Teoh, A. Leung, K. Ludlow, I. Cohen were supported in part by a National Institute of Neurological Disorders and Stroke, NIH grant: 5R01NS116595. H.K. Teoh, D. Biswas, A. Leung, K. Ludlow, N. J. Cowan and I. Cohen were supported by a National Institute of Neurological Disorders and Stroke, NIH grant: 5U01NS131438. S. Whitehead was supported by Della Martin Postdoctoral Fellowship. M. Dickinson was supported by a National Institute of Neurological Disorders and Stroke, NIH grant: 5R01NS136988.

## Supplementary Note 1: Supplementary Information

### A. FANC Dataset: Premotor neurons from haltere tract innervating wing motoneurons

**Figure S1.**
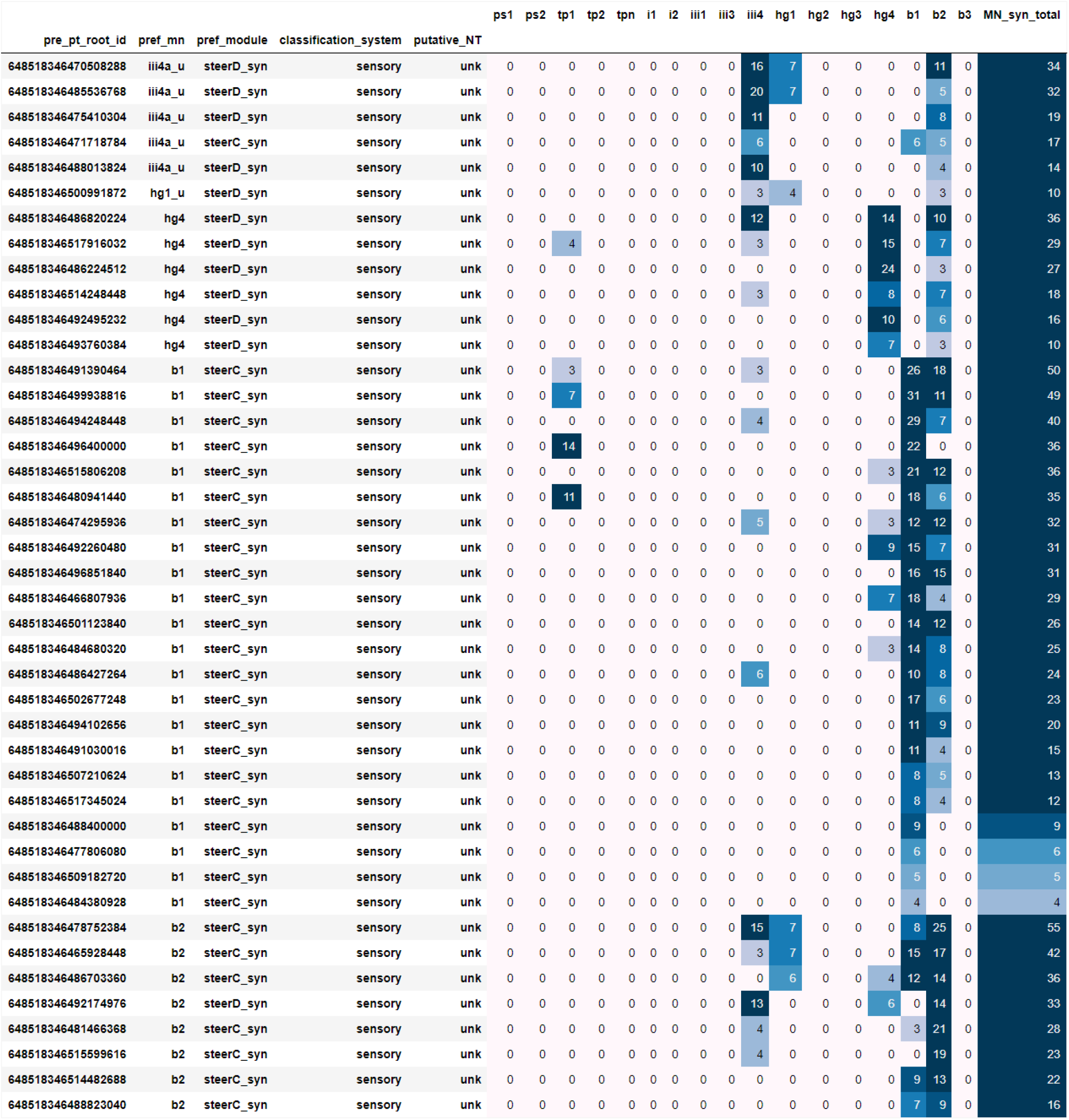
Premotor neurons from haltere tract innervating b1, b2 motoneurons

**Figure S2.**
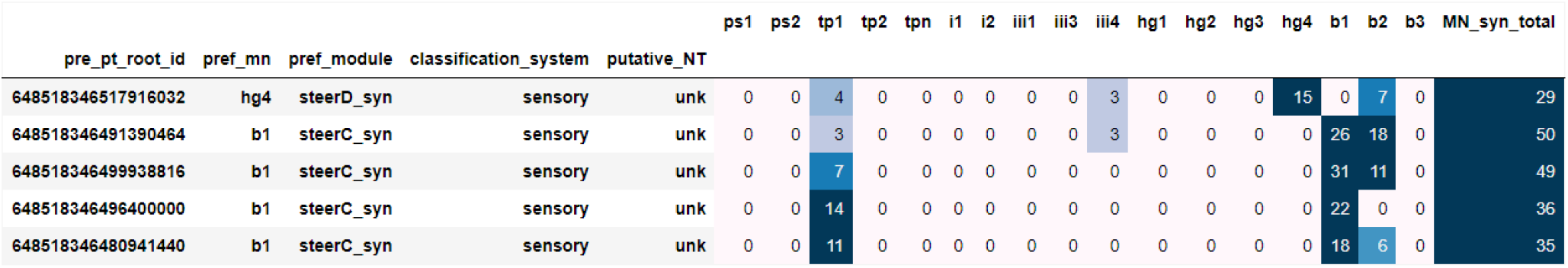
Premotor neurons from haltere tract innervating tp1 motoneuron

**Figure S3.**
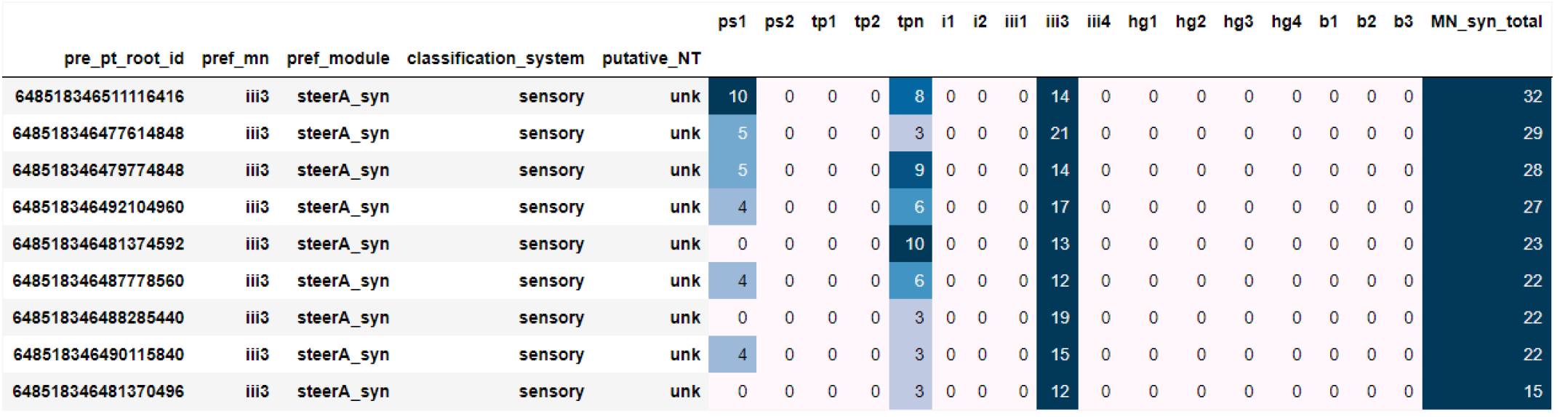
Premotor neurons from haltere tract innervating tpn motoneuron

**Figure S4.**
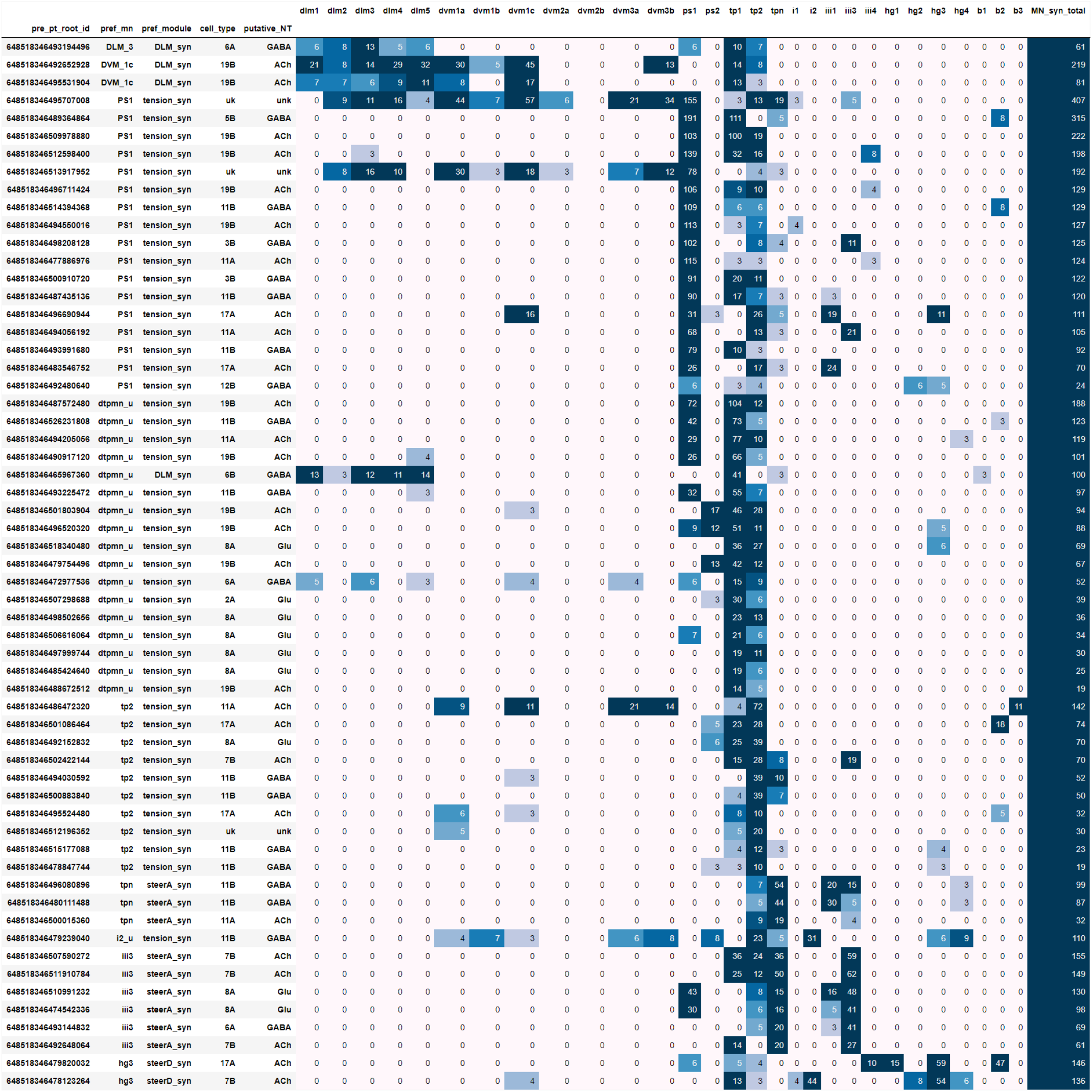
Local premotor neurons innervating tp motoneurons

## Supplementary Note 2: Maximum intensity projection of split driver lines

**Figure S5.**
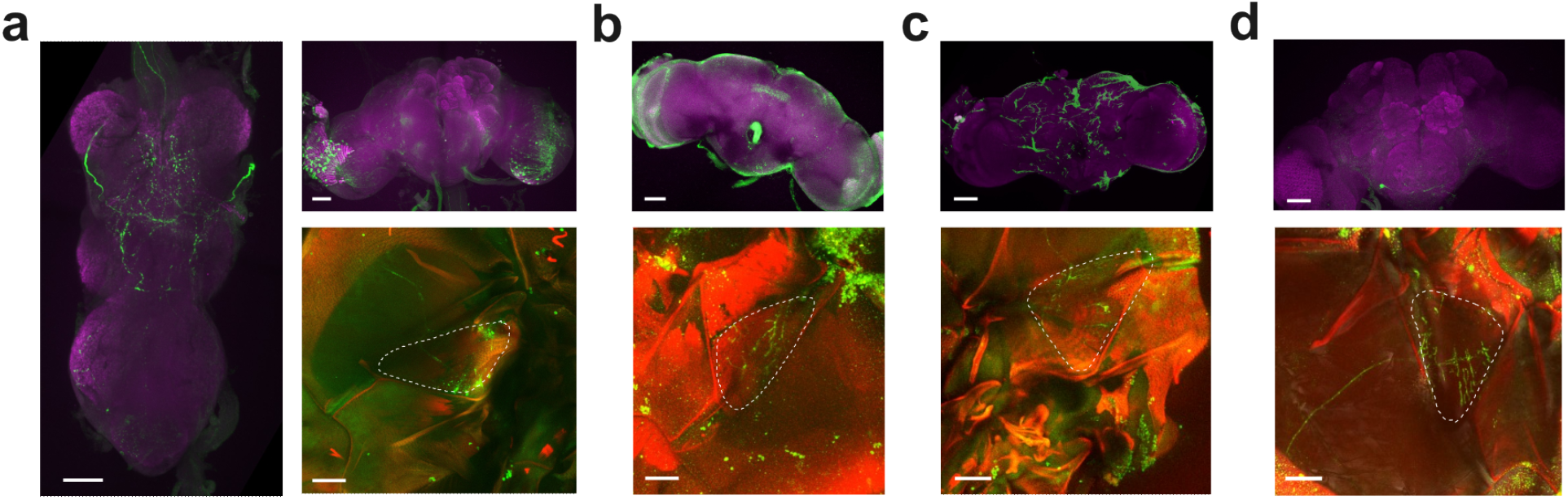
Maximum intensity projection (MIP) images of the split driver lines tp1,tp1-SG,tp2 and tpN. (a) Left - A Maximum intensity projection (MIP) VNC image from a tp1-SG > CsChrimson fly. Green corresponds to mVenus, purple to DNCad (neuropil). Top right-MIP brain image from a tp1-SG > CsChrimson fly. Bottom right - Phalloidin-stained thoracic hemisection from a tp1-SG > CsChrimson fly showing wing musculature, with both phalloidin (red) and GFP (green) expression innervating the tp1 muscle (outlined with white dashed lines). (b) Top - same as (a) top right but with tp1-GAL4 > CsChrimson. Bottom - same as (a) bottom left except with tp1-GAL4 > CsChrimson. (c) Top - same as (a) top right but with tp2-GAL4 > CsChrimson. Bottom - same as (a) bottom left except with tp2-GAL4 > CsChrimson. (d) Top - same as (a) top right but with tpN-GAL4 > CsChrimson. Bottom - same as (a) bottom left except with tpN-GAL4 > CsChrimson. All scale bars 50 *µ*m. See Table 1 for full fly genotypes

## Supplementary Note 3: Lift Maximization Model Predictions

**Figure S6.**
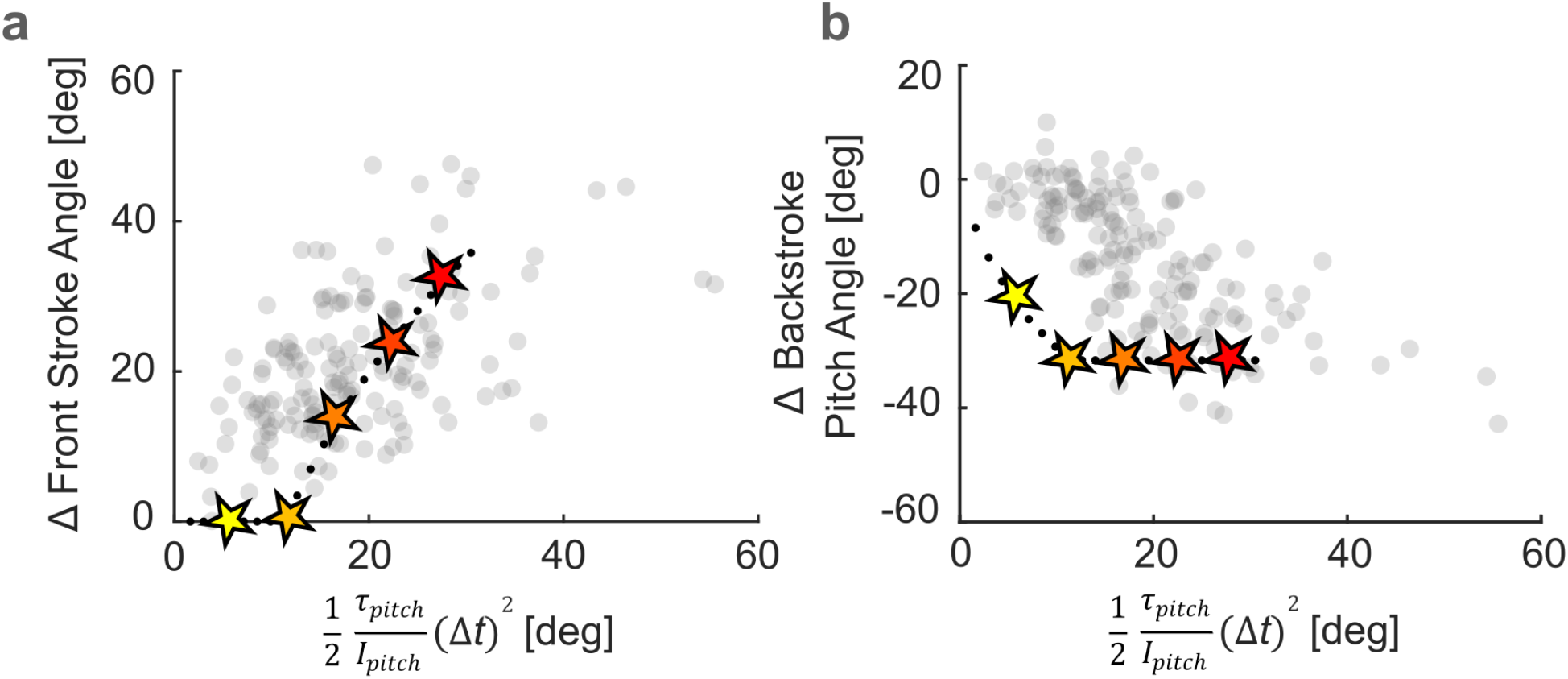
Lift maximization model prediction. (a) The backstroke wing pitch response and (b) front stroke response generated from the lift maximization model in Fig. 2 (n). The grey dots are experimental data. The dotted lines represent the model prediction, and the star symbols correspond to the example contours discussed in the main text.

## Supplementary Note 4: Tp muscles optogenetic activation and inhibition analysis

**Table 2.** Statistical results for Fig. 3. Summary of nonparametric group comparisons for wing kinematics and behavioral responses under tp1 inhibition and activation. *H* refers to the Kruskal–Wallis test statistic; *p* is the corresponding uncorrected significance value. *η*^2^ indicates the effect size estimate for Kruskal–Wallis tests, and *h* denotes Cohen’s *h* for pairwise proportion comparisons. Post hoc *p*-values are from multiple comparisons test with Dunnett correction (for panels a–d, f–i), or Bonferroni-corrected pairwise proportion tests (for panels e, j). See Supplementary Table 3 for effect size interpretations.

| Panel | Measure | Comparison | $H$ | $p$ | $\eta^2 / h$ | Post hoc $p$ |
| --- | --- | --- | --- | --- | --- | --- |
| (a, b) | Wing pitch (Small perturbation) | control vs tp1<br>control vs tp1-alt | 0.97 | 0.616 | 0 | 0.991<br>0.676 |
|  | Wing stroke (Small perturbation) | control vs tp1<br>control vs tp1-alt | 4.76 | 0.093 | 0.084 | 0.999<br>0.132 |
| (c, d) | Wing pitch (Big perturbation) | control vs tp1<br>control vs tp1-alt | 9.58 | 0.008 | 0.100 | 0.026<br>0.008 |
|  | Wing stroke (Big perturbation) | control vs tp1<br>control vs tp1-alt | 4.05 | 0.132 | 0.027 | 0.261<br>0.094 |
| (e) | Correction (>50%) | control vs tp1<br>control vs tp1-alt | — | 0.0075<br><0.001 | $h = -0.755$<br>$h = -1.446$ | —<br>— |
| (f, g) | Wing pitch (Small perturbation) | control vs tp1<br>control vs tp1-alt | 9.41 | 0.009 | 0.151 | 0.008<br>0.023 |
|  | Wing stroke (Small perturbation) | control vs tp1<br>control vs tp1-alt | 9.50 | 0.009 | 0.153 | 0.101<br>0.004 |
| (h, i) | Wing pitch (Big perturbation) | control vs tp1<br>control vs tp1-alt | 0.12 | 0.944 | 0 | 0.926<br>0.951 |
|  | Wing stroke (Big perturbation) | control vs tp1<br>control vs tp1-alt | 2.71 | 0.258 | 0.012 | 0.958<br>0.223 |
| (j) | Over-correction (>10%) | control vs tp1<br>control vs tp1-alt | — | 0.0030<br>0.0200 | $h = 1.160$<br>$h = 0.890$ | —<br>— |

**Table 3.** Interpretation of Cohen’s *h* effect sizes used in Fig. 3.

| Panel | Comparison | Cohen's $h$ | Interpretation |
| --- | --- | --- | --- |
| (e) | control vs tp1 | −0.755 | Large |
|  | control vs tp1-alt | −1.446 | Very large |
| (j) | control vs tp1 | 1.160 | Very large |
|  | control vs tp1-alt | 0.890 | Large |

**Figure S7.**
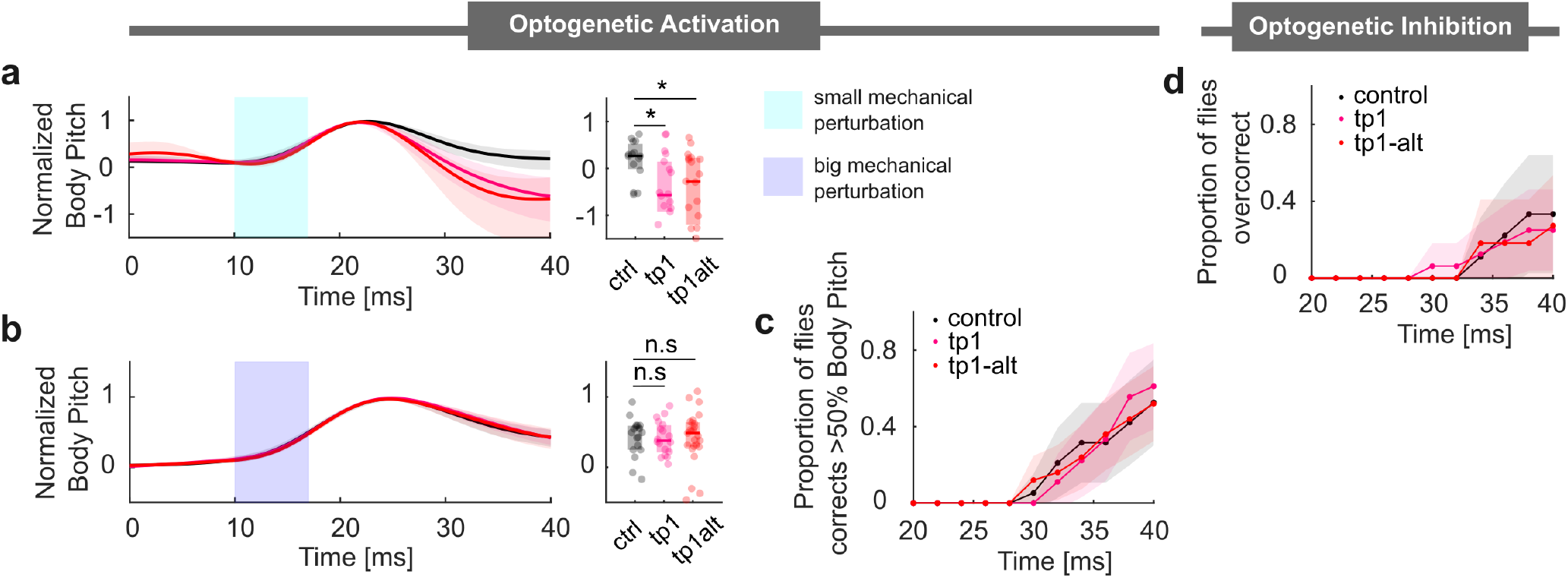
Additional tp1 activation and inhibition plots. (a and b) Left: The normalized body pitch angle of control and tp1-activated flies along the pitch axis during a correction maneuver for small and big pitch perturbation. Right: Normalized body pitch angle at time 40ms. (c) The proportion of flies correcting more than 50% of control flies body deflection amplitude at time 40ms. (d) The proportion of flies over-correcting more than 10% of control flies body deflection amplitude at time 40ms. Statistical significance for (a)-(b) is determined via the Kruskal-Wallis test with Dunn’s post hoc multiple comparisons. Statistical significance for (c) and (d) is determined via the Proportional test with Bonferroni correction(***, *p <*.001; **, *p <*.01; *, *p <*.05). Optogenetic inhibition small perturbation: control *n*=12, tp1 *n*= 24, tp1-alt *n*=15. Optogenetic activation small perturbation: control, *n*=16, tp1 *n*= 17, tp1-alt *n*= 11. Optogenetic activation big perturbation: control, *n*=19, tp1 *n*= 16, tp1-alt *n*= 11).

**Figure S8.**
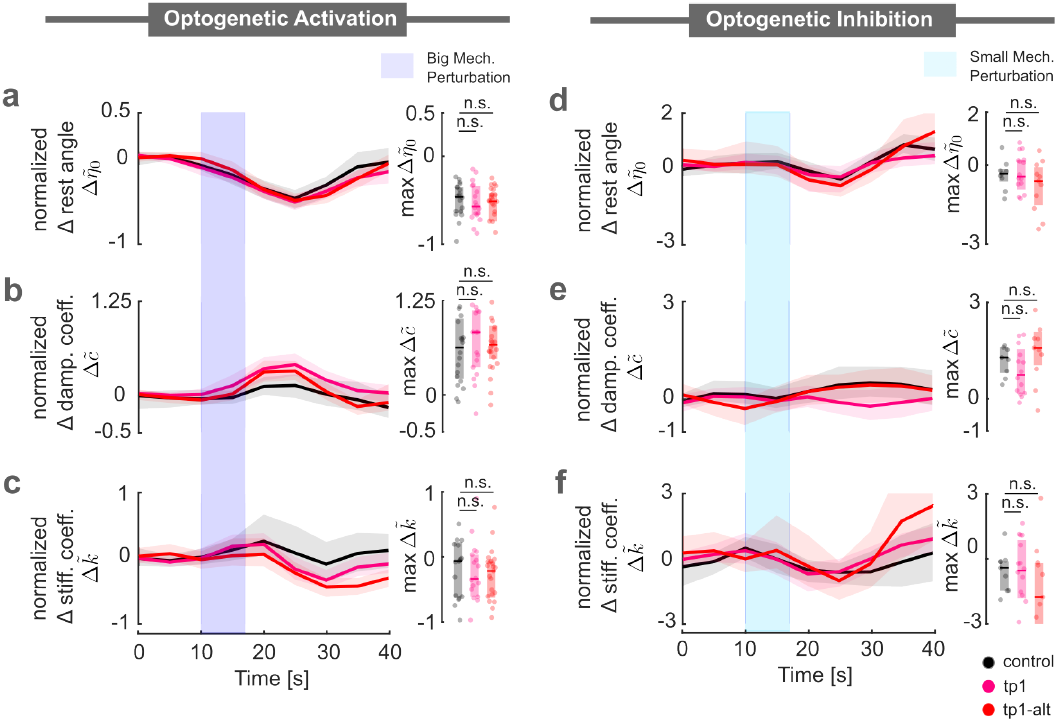
Additional tp1 torsional spring fit plots. (a) Left: Normalized change in the torsional spring rest angle: Comparison of control and tp1-activated flies in response to large pitch perturbations. Right: Maximum change in normalized torsional spring rest angle. (b) Left: Normalized change in the torsional spring damping coefficient: Comparison of control and tp1-activated flies in response to large pitch perturbations. Right: Maximum change in normalized torsional spring damping coefficient. (c) Left: Normalized change in the torsional spring stiffness constant: Comparison of control and tp1-activated flies in response to large pitch perturbations. Right: Maximum change in normalized torsional spring stiffness constant. (d) Left: Normalized change in the torsional spring rest angle: Comparison of control and tp1-inhibited flies in response to small pitch perturbations. Right: Maximum change in normalized torsional spring rest angle. (e) Left: Normalized change in the torsional spring damping coefficient: Comparison of control and tp1-inhibited flies in response to small pitch perturbations. Right: Maximum change in normalized torsional spring damping coefficient. (f) Left: Normalized change in the torsional spring stiffness constant: Comparison of control and tp1-inhibited flies in response to small pitch perturbations. Right: Maximum change in normalized torsional spring stiffness constant. Statistical significance for (a)-(f) is determined via the Kruskal-Wallis test with Dunn’s post hoc multiple comparisons. (***, *p <*.001; **, *p <*.01; *, *p <*.05). Optogenetic activation big perturbation: control, *n*=19, tp1 *n*= 16, tp1-alt *n*= 11. Optogenetic inhibition small perturbation: control *n*=12, tp1 *n*= 24, tp1-alt *n*= 15.

**Figure S9.**
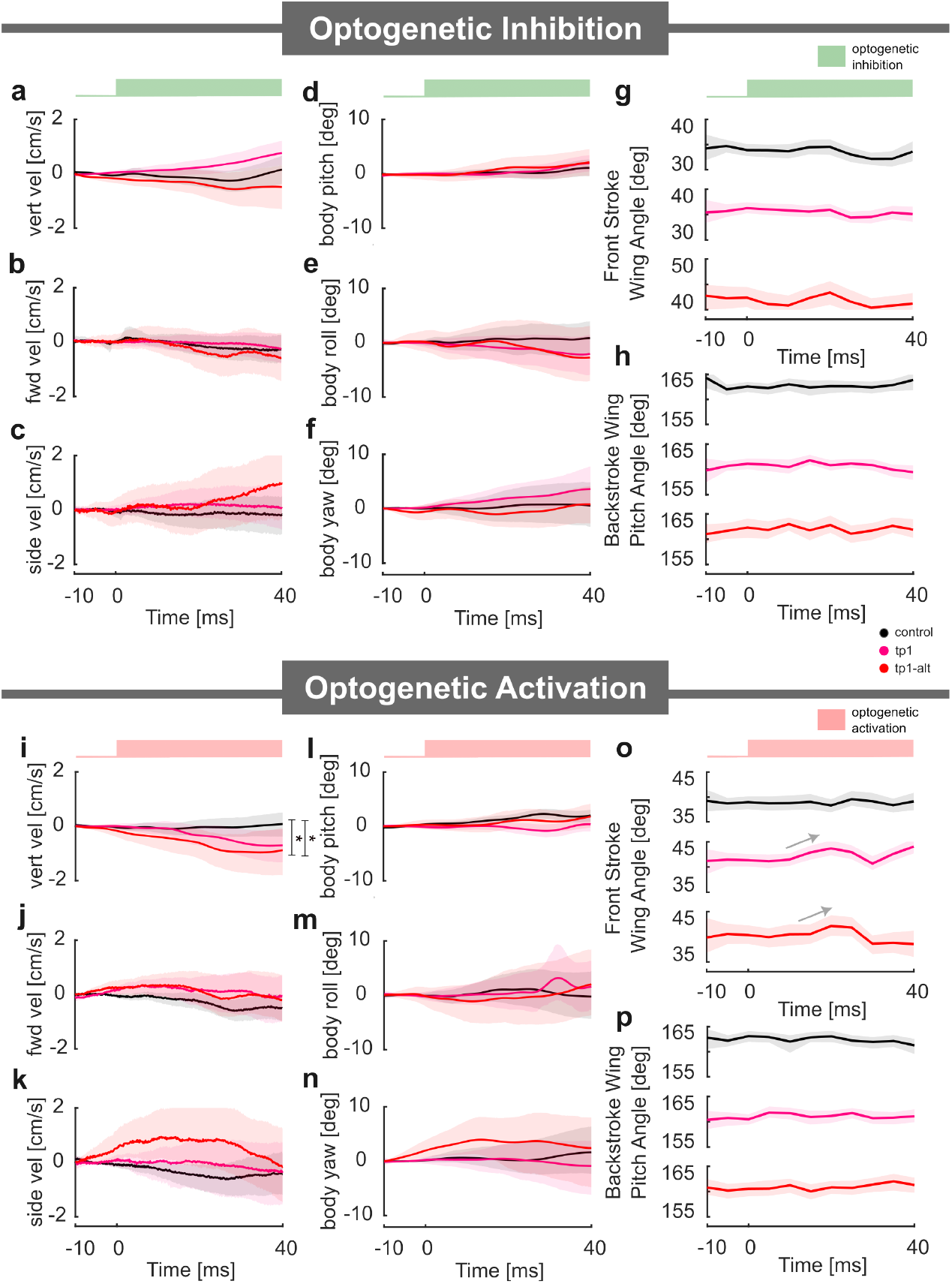
tp1 inhibition and activation data. (First column: a, b, c, i, j and k) Translational velocities (Vertical, forward, and side) of control and tp1-inhibited/activated flies. (Second column: d, e, f, l, m and n) Angular orientation (body pitch, body roll, and body yaw) of control and tp1-inhibited/activated flies. (Third column: g, h, o, and p) Front stroke wing angle and backstroke pitch angle of control and tp1-inhibited/activated flies. In (i) tp1-activated flies show a downward drift compared to controls. A linear slope was estimated for each time series within the groups to quantify this change. Distributionally, the slope for tp1-activated flies exhibits a heavier tail. (Proportion test at the 10% percentile of control, *p <*.01). The observed vertical velocity drift in tp1-activated flies can be attributed to a slight decrease in the front stroke wing angle (panel o), which leads to reduced lift generation. Sample sizes for optogenetic inhibition: control *n*= 94, tp1 *n*= 93 and tp1-alt *n*= 46. Sample sizes for optogenetic activation: control, *n*=93, tp1 *n*= 93 and tp1-alt *n*= 57

**Figure S10.**
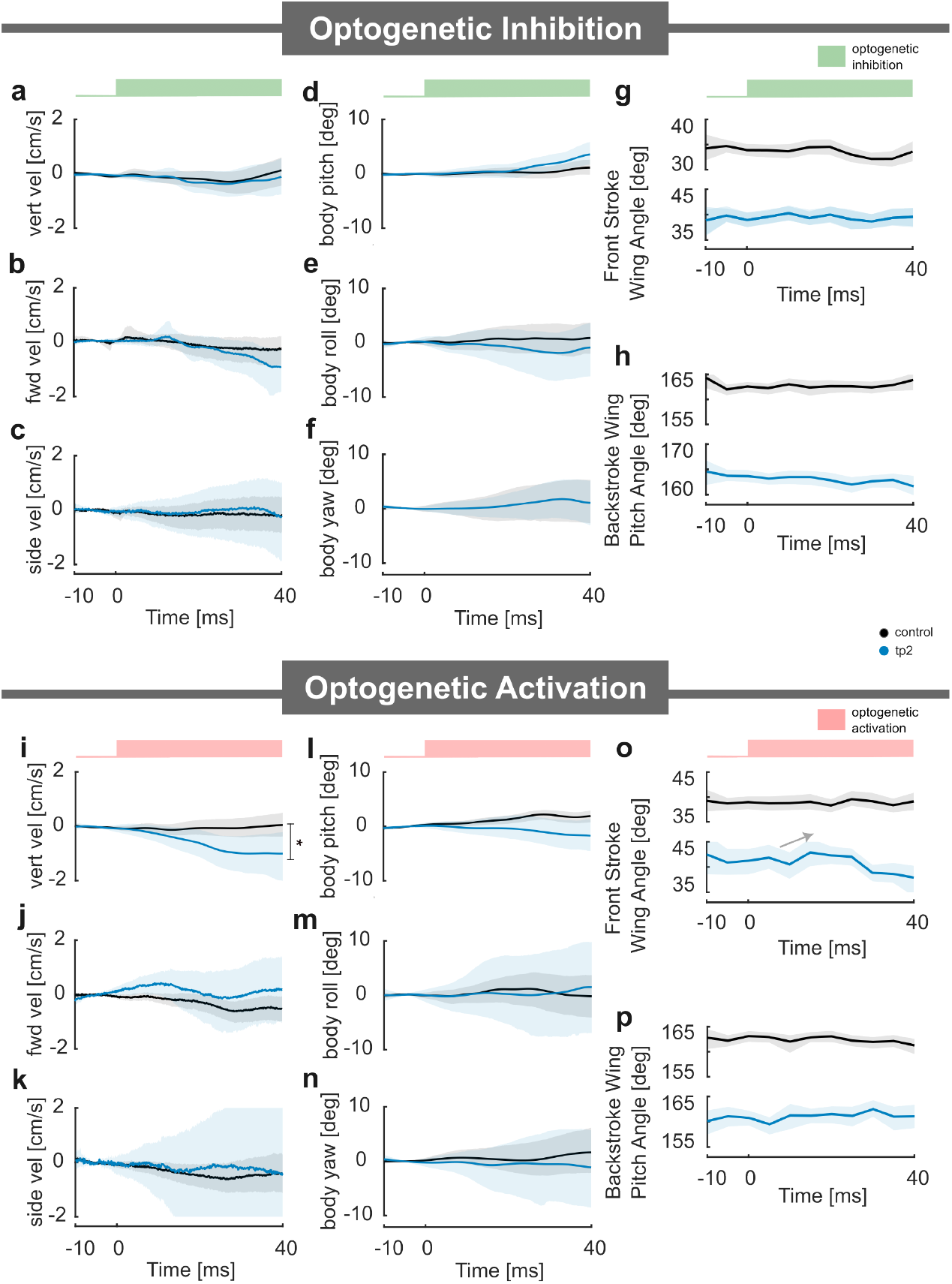
tp2 free flight inhibition and activation data. (First column: a, b, c, i, j and k) Translational velocities (Vertical, forward, and side) of control and tp2-inhibited/activated flies. (Second column: d, e, f, l, m and n) Angular orientation (body pitch, body roll, and body yaw) of control and tp2-inhibited/activated flies. (Third column: g, h, o, and p) Front stroke wing angle and backstroke pitch angle of control and tp1-inhibited/activated flies. In (i) tp2-activated flies show a downward drift compared to controls. A linear slope was estimated for each time series within the groups to quantify this change. Distributionally, the slope for tp2-activated flies exhibits a heavier tail. (Proportion test at the 10% percentile of control, *p <*.01). The observed vertical velocity drift in tp1-activated flies can be attributed to a slight decrease in the front stroke wing angle (panel o), which leads to reduced lift generation Sample sizes for optogenetic inhibition: control *n*= 94, tp2 *n*= 59. Sample sizes for optogenetic activation: control, *n*=93, tp2 *n*= 39

**Figure S11.**
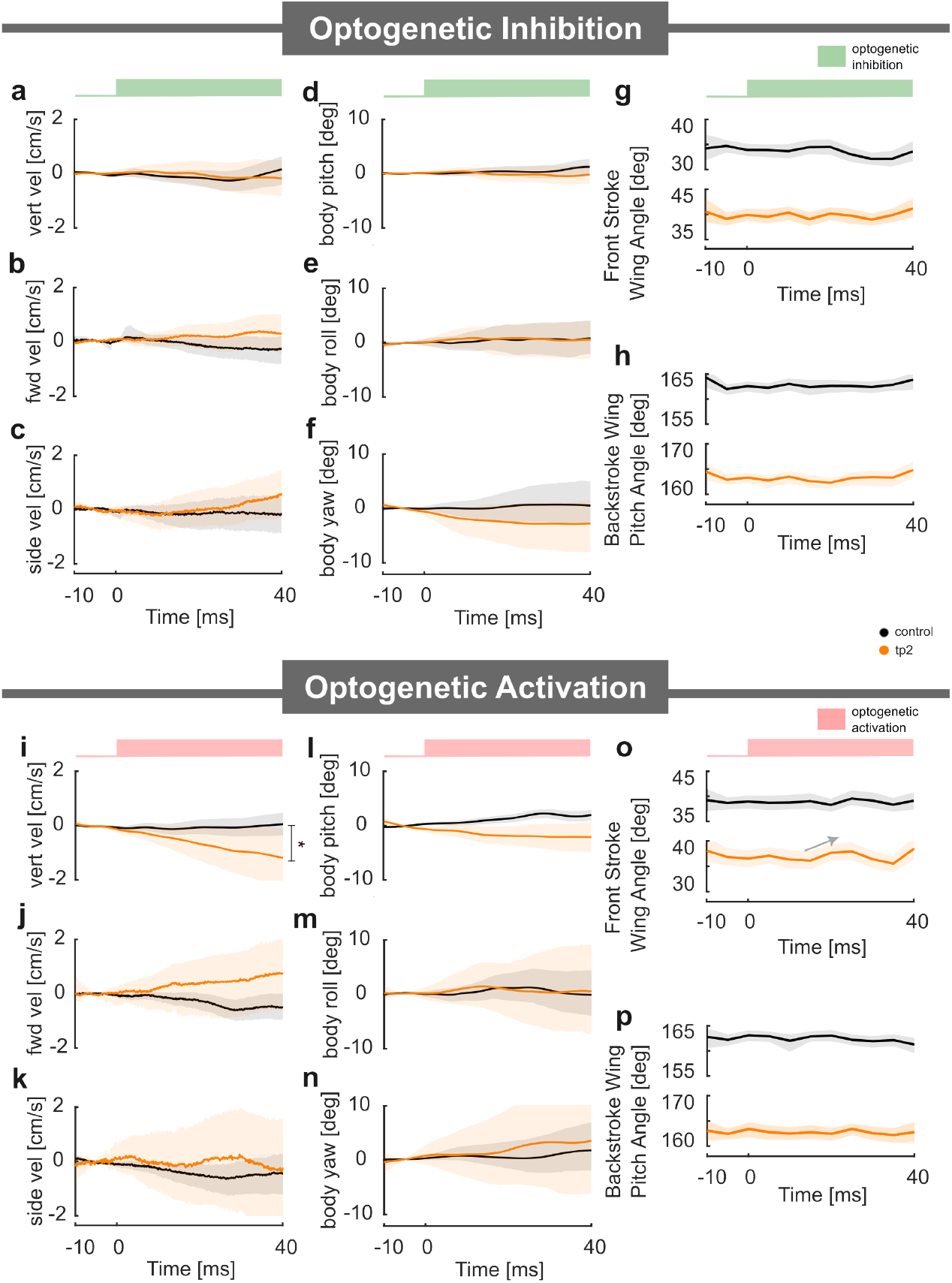
tpN free flight inhibition and activation data. (First column: a, b, c, i, j and k) Translational velocities (Vertical, forward, and side) of control and tpN-inhibited/activated flies. (Second column: d, e, f, l, m and n) Angular orientation (body pitch, body roll, and body yaw) of control and tpN-inhibited/activated flies. (Third column: g, h, o, and p) Front stroke wing angle and backstroke pitch angle of control and tpN-inhibited/activated flies. In (i) tpN-activated flies show a downward drift compared to controls. A linear slope was estimated for each time series within the groups to quantify this change. Distributionally, the slope for tpN-activated flies exhibits a heavier tail. (Proportion test at the 10% percentile of control, *p <*.01). The observed vertical velocity drift in tpN-activated flies can be attributed to a slight decrease in the front stroke wing angle (panel o), which leads to reduced lift generation Sample sizes for optogenetic inhibition: control *n*= 94, tpN *n*= 54. Sample sizes for optogenetic activation: control, *n*=93, tpN *n*= 27

**Figure S12.**
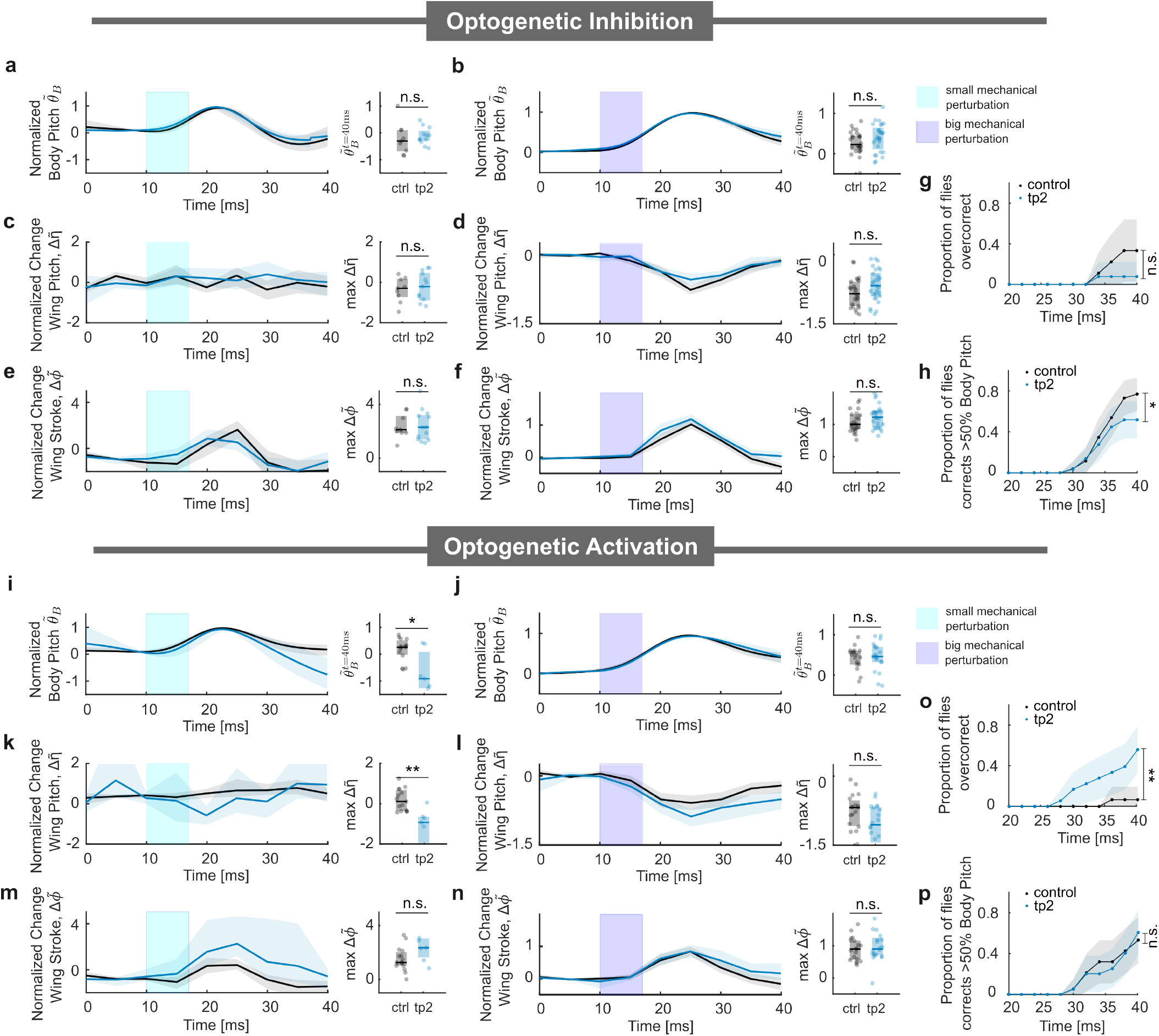
tp2 inhibition and activation data. (a and b) Left: Normalized body pitch angle of control and tp2-inhibited flies along the pitch axis during a correction maneuver for small and big pitch perturbations. Right: Normalized body pitch angle at time 40ms. (c and d) Left: Normalized change in backstroke wing pitch response: Comparison of control and tp2-inhibited flies in response to small and big pitch perturbations. Right: Maximum change in normalized wing pitch response. (e and f) Left: Normalized change in front stroke response: Comparison of control and tp2-inhibited flies in response to small and big pitch perturbations. Right: Maximum change in normalized wing stroke response (g) The proportion of flies over-correcting more than 10% of control flies body deflection amplitude at time 40ms. (h) The proportion of flies that correct more than 50 % body deflection amplitude during a big pitch perturbation as a function of time. (i and j) Left: Normalized body pitch angle of control and tp2-activated flies along the pitch axis during a correction maneuver for small and big pitch perturbations. Right: Normalized body pitch angle at time 40ms. (k and l) Normalized change in backstroke wing pitch response: Comparison of control and tp2-activated flies in response to small and big pitch perturbations. Right: Maximum change in normalized wing pitch response. (m and n) Left: Normalized change in front stroke response: Comparison of control and tp2-activated flies in response to small and large pitch perturbations. Right: Maximum change in normalized wing stroke response. (o) The proportion of flies over-correcting more than 10% of control flies body deflection amplitude at time 40ms. (p) The proportion of flies that correct more than 50 % body deflection amplitude during a big pitch perturbation as a function of time. Optogenetic inhibition small perturbation: control *n*=9, tp2 *n*= 13. Optogenetic inhibition big perturbation: control *n*=26, tp2 *n*= 29. Statistical significance for (a)-(f) and (i)-(n) are determined via the Kruskal-Wallis test with Dunn’s post hoc multiple comparisons. Statistical significance for (g), (h), (o), and (p) are determined via the Proportional test with Bonferroni correction(***, *p <*.001; **, *p <*.01; *, *p <*.05). Optogenetic activation small perturbation: control, *n*=15, tp2 *n*= 7. Optogenetic activation big perturbation: control, *n*=19, tp2 *n*= 20

**Figure S13.**
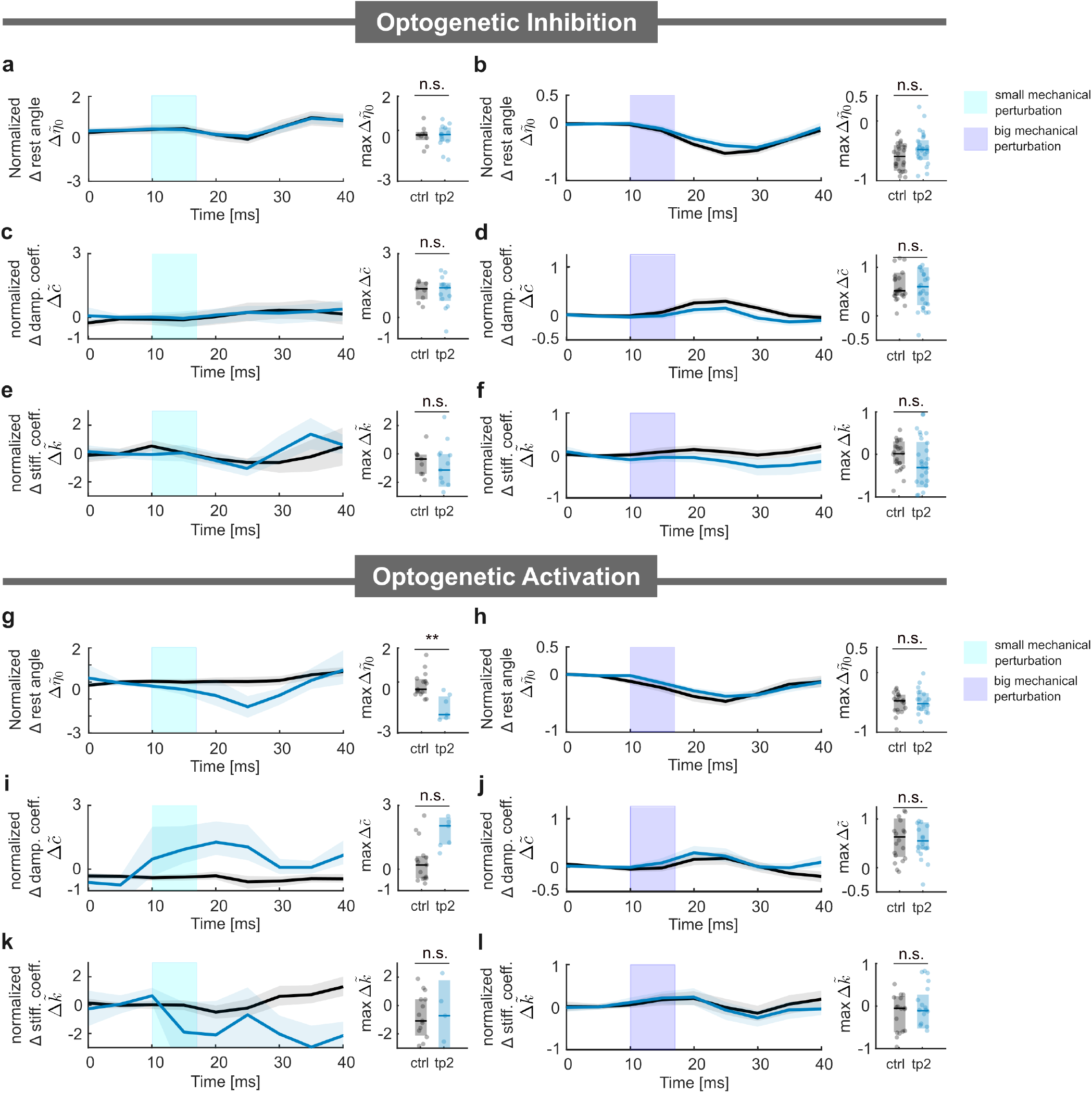
tp2 torsional spring model fit. (a,b) Left: Normalized change in the torsional spring rest angle: Comparison of control and tp2-inhibited flies in response to small and large pitch perturbations. Right: Maximum change in normalized torsional spring rest angle. (c,d) Left: Normalized change in the torsional spring damping factor: Comparison of control and tp2-inhibited flies in response to small and large pitch perturbations. Right: Maximum change in normalized torsional spring damping factor. (e,f) Left: Normalized change in the torsional spring stiffness constant: Comparison of control and tp2-inhibited flies in response to small and large pitch perturbations. Right: Maximum change in normalized torsional spring stiffness constant. (g,h) Left: Normalized change in the torsional spring rest angle: Comparison of control and tp2-activated flies in response to small and large pitch perturbations. Right: Maximum change in normalized torsional spring rest angle. (i,j) Left: Normalized change in the torsional spring damping factor: Comparison of control and tp2-activated flies in response to small and large pitch perturbations. Right: Maximum change in normalized torsional spring damping factor. (k,l) Left: Normalized change in the torsional spring stiffness constant: Comparison of control and tp2-activated flies in response to small and large pitch perturbations. Right: Maximum change in normalized torsional spring stiffness constant. Statistical significance for (a)-(l) is determined via the Kruskal-Wallis test with Dunn’s post hoc multiple comparisons. (***, *p <*.001; **, *p <*.01; *, *p <*.05). In (g) we observed a significant change in rest angle relative of tp2 activated flies to control (Kruskal-Wallis test, *p <*.001). In panels (i) and (k), the damping factor and stiffness constant estimations exhibit high variability, even before mechanical perturbation, likely due to the small sample size. Optogenetic inhibition small perturbation: control *n*=9, tp2 *n*= 13. Optogenetic inhibition big perturbation: control *n*=26, tp2 *n*= 29. Optogenetic activation small perturbation: control, *n*=15, tp2 *n*= 6. Optogenetic activation big perturbation: control, *n*=19, tp2 *n*= 19

**Figure S14.**
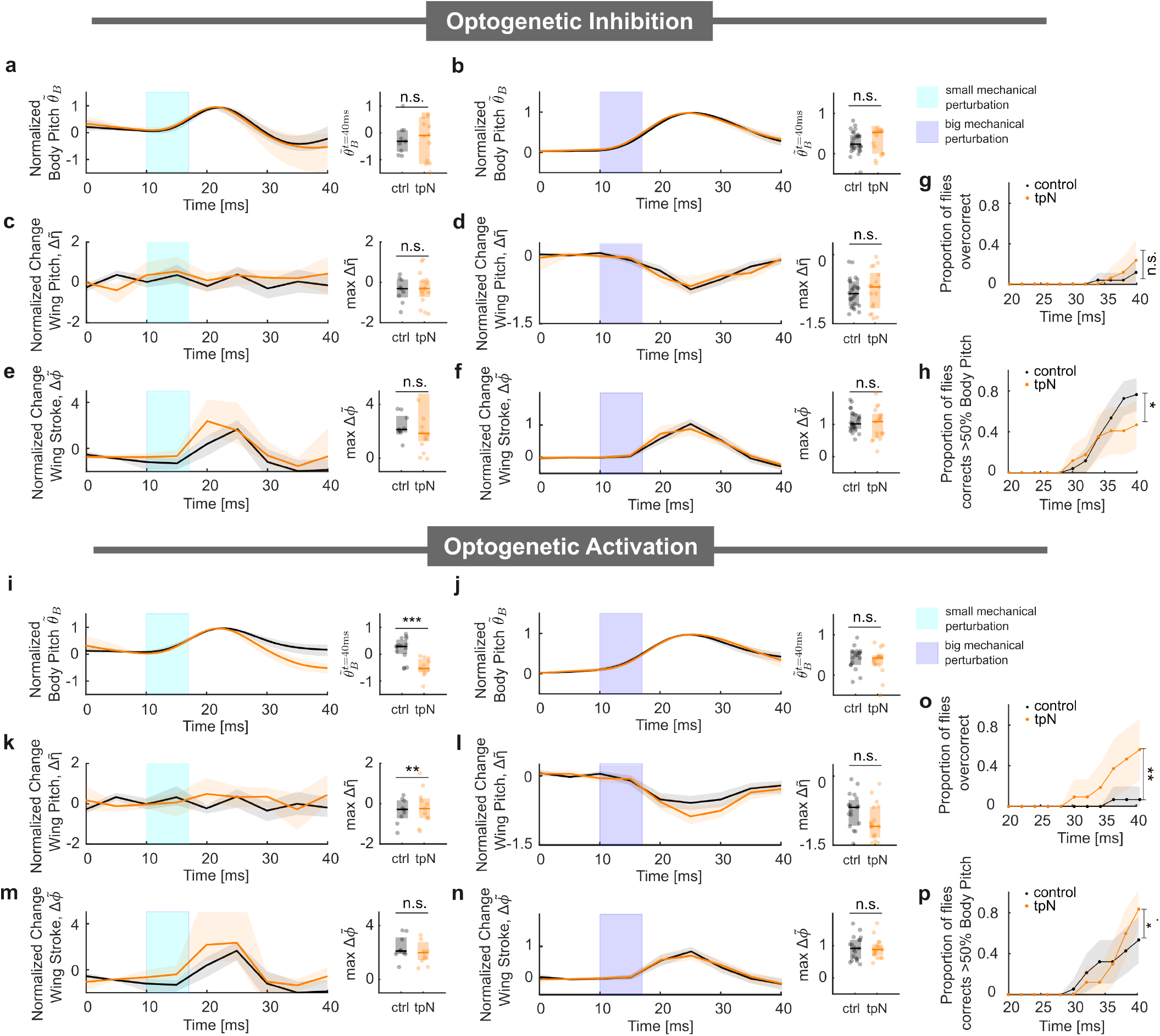
tpN inhibition and activation data. (a and b) Left: Normalized body pitch angle of control and tpN-inhibited flies along the pitch axis during a correction maneuver for small and big pitch perturbations. Right: Normalized body pitch angle at time 40ms. (c and d) Left: Normalized change in backstroke wing pitch response: Comparison of control and tpN-inhibited flies in response to small and big pitch perturbations. Right: Maximum change in normalized wing pitch response. (e and f) Left: Normalized change in front stroke response: Comparison of control and tpN-inhibited flies in response to small and big pitch perturbations. Right: Maximum change in normalized wing stroke response (g) The proportion of flies over-correcting more than 10% of control flies body deflection amplitude at time 40ms. (h) The proportion of flies that correct more than 50 % body deflection amplitude during a big pitch perturbation as a function of time. (i and j) Left: Normalized body pitch angle of control and tpN-activated flies along the pitch axis during a correction maneuver for small and big pitch perturbations. Right: Normalized body pitch angle at time 40ms. (k and l) Normalized change in backstroke wing pitch response: Comparison of control and tpN-activated flies in response to small and big pitch perturbations. Right: Maximum change in normalized wing pitch response. (m and n) Left: Normalized change in front stroke response: Comparison of control and tpN-activated flies in response to small and large pitch perturbations. Right: Maximum change in normalized wing stroke response. (o) The proportion of flies over-correcting more than 10% of control flies body deflection amplitude at time 40ms. (p) The proportion of flies that correct more than 50 % body deflection amplitude during a big pitch perturbation as a function of time. Statistical significance for (a)-(f) and (i)-(n) are determined via the Kruskal-Wallis test with Dunn’s post hoc multiple comparisons. Statistical significance for (g), (h), (o), and (p) are determined via the Proportional test with Bonferroni correction(***, *p <*.001; **, *p <*.01; *, *p <*.05). Optogenetic inhibition small perturbation: control *n*=9, tpN *n*= 16. Optogenetic inhibition big perturbation: control *n*=26, tpN *n*= 17. Optogenetic activation small perturbation: control, *n*=15, tpN *n*= 11. Optogenetic activation big perturbation: control, *n*=19, tpN *n*= 17

**Figure S15.**
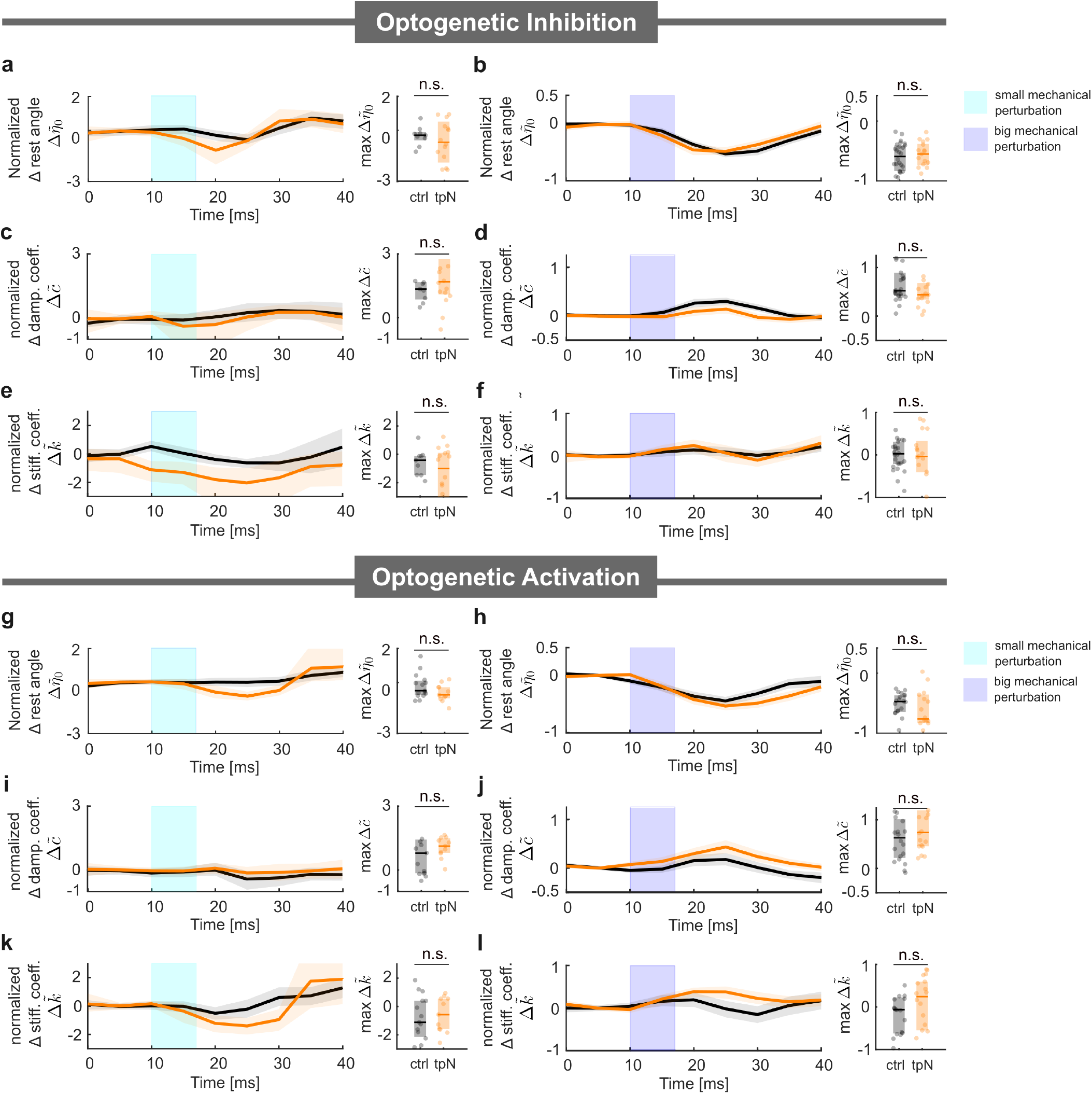
tpN torsional spring model fit. (a,b) Left: Normalized change in the torsional spring rest angle: Comparison of control and tpN-inhibited flies in response to small and large pitch perturbations. Right: Maximum change in normalized torsional spring rest angle. (c,d) Left: Normalized change in the torsional spring damping factor: Comparison of control and tpN-inhibited flies in response to small and large pitch perturbations. Right: Maximum change in normalized torsional spring damping factor. (e,f) Left: Normalized change in the torsional spring stiffness constant: Comparison of control and tpN-inhibited flies in response to small and large pitch perturbations. Right: Maximum change in normalized torsional spring stiffness constant. (g,h) Left: Normalized change in the torsional spring rest angle: Comparison of control and tpN-activated flies in response to small and large pitch perturbations. Right: Maximum change in normalized torsional spring rest angle. (i,j) Left: Normalized change in the torsional spring damping factor: Comparison of control and tpN-activated flies in response to small and large pitch perturbations. Right: Maximum change in normalized torsional spring damping factor. (k,l) Left: Normalized change in the torsional spring stiffness constant: Comparison of control and tpN-activated flies in response to small and large pitch perturbations. Right: Maximum change in normalized torsional spring stiffness constant. Statistical significance for (a)-(l) is determined via the Kruskal-Wallis test with Dunn’s post hoc multiple comparisons. (***, *p <*.001; **, *p <*.01; *, *p <*.05). Optogenetic inhibition small perturbation: control *n*=9, tpN *n*= 16. Optogenetic inhibition big perturbation: control *n*=26, tpN *n*= 17. Optogenetic activation small perturbation: control, *n*=15, tpN *n*= 11. Optogenetic activation big perturbation: control, *n*=19, tpN *n*= 17

## Supplementary Note 5: Additional control theory analysis

**Figure S16.**
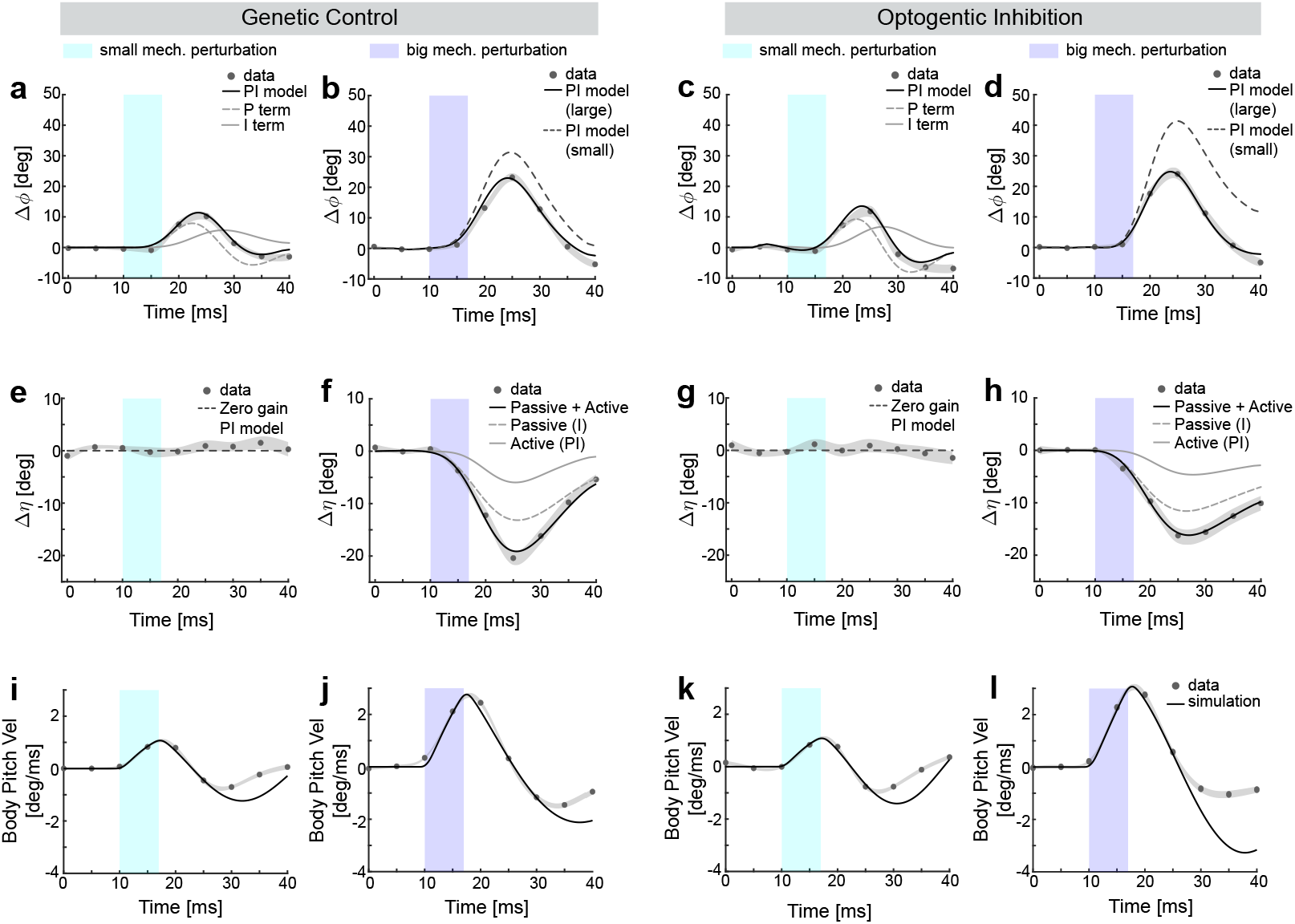
Control theoretic model fits for genetic control and tp1 inhibition data. (a–d) Change in wing stroke angle over time for simulated flies (black lines, PI controller) fitted to the experimental data (black circles representing the population average, with the gray shaded region indicating the SEM) for small (a,c) and large (b,d) mechanical perturbation in the control (a, b) and tp1 inhibited (c, d) groups. Gray dashed and solid lines in (a,c) denote the contributions from the P term (proportional), and the I term (integral), respectively. The gray dashed line in (b,d) denotes the PI controller model fit with parameters from small perturbation. (e–h) Change in wing pitch angle over time for simulated flies (black lines) fitted to the experimental data (black circles representing the population average, with the gray shaded region indicating the SEM) for small (e,g) and big (f,h) perturbation in the control (e, f) and tp1 inhibited (g, h) groups. The dashed lines in (e,g) denote the zero gain PI controller model fit to the small perturbation data. The solid black line, dashed gray line, and solid gray line in (f,h) represent the combined passive and active controller model fit, the individual passive model fit, and the active model fit, respectively. (i–l) Change in body pitch velocity over time for simulated flies. The colors are same as in (a–h). Small perturbation: control (control) *n*=28, optogenetic inhibition (tp1 inh) *n*=27. Big perturbation: control *n*=61, tp1 inh *n*= 53.

**Table 4.** Fitted controller parameter values with mean and standard deviation.

|  | Stroke Controller |  |  |  | Pitch Controller Passive |  | Pitch Controller Active |  |  |  |
| --- | --- | --- | --- | --- | --- | --- | --- | --- | --- | --- |
| | $K_p$ (/ms) | | $K_i$ [deg/deg] | | $K_i$ [deg/deg] | | $K_p$ (/ms) | | $K_i$ [(deg/deg)] | |
|  | Small | Large | Small | Large | Small | Large | Small | Large | Small | Large |
| Control | 7.5794±0.5503 | 6.2930±0.3821 | 0.6931±0.1028 | 0.3984±0.0472 | 0 | -0.5±0.0998 | 0 | -1.2±0.1499 | 0 | -0.1616±0.0167 |
| tp1 act | 7.0000±1.0433 | 5.9261±0.1831 | 1.4±0.2250 | 0.5876±0.0266 | -0.6280±0.0373 | -0.65±0.0585 | -0.8±0.2635 | -1±0.3881 | -0.0840±0.0562 | -0.2489±0.0686 |
| tp1 inh | 8.6030±0.8889 | 7.0803±0.2982 | 0.7±0.1986 | 0.2299±0.0344 | 0 | -0.4±0.0223 | 0 | -0.5099±0.0738 | 0 | -0.1472±0.0068 |
| tp2 act | 5.0806±0.7564 | 4.90173±0.3468 | 0.9286±0.1320 | 0.7478±0.0560 | -0.5455±0.2541 | -0.85±0.0393 | -1±1.5922 | -0.9±0.06426 | -0.0840±0.2705 | -0.1±0.0945 |
| tp2 inh | 7.5801±0.3242 | 6.9967±0.2289 | 0.9178±0.0643 | 0.9379±0.0327 | 0 | -0.3±0.0866 | 0 | -0.6422±0.2795 | 0 | -0.3497±0.0372 |
| tpn act | 7.9081±0.2926 | 5.1694±0.1242 | 0.4602±0.0384 | 0.5476±0.0166 | 0±0.0569 | -0.85±0.0545 | 0±0.4495 | -1±0.4981 | 0±0.0684 | -0.1744±0.0755 |
| tpn inh | 7.5051±0.7256 | 5.5576±0.4266 | 0.8875±0.1515 | 0.7037±0.0702 | 0 | -0.4±0.1368 | 0 | -0.3092±0.1338 | 0 | -0.2676±0.0164 |

## Supplementary Note 6: Quasi-steady aerodynamic calculation parameter values

**Table 5.**
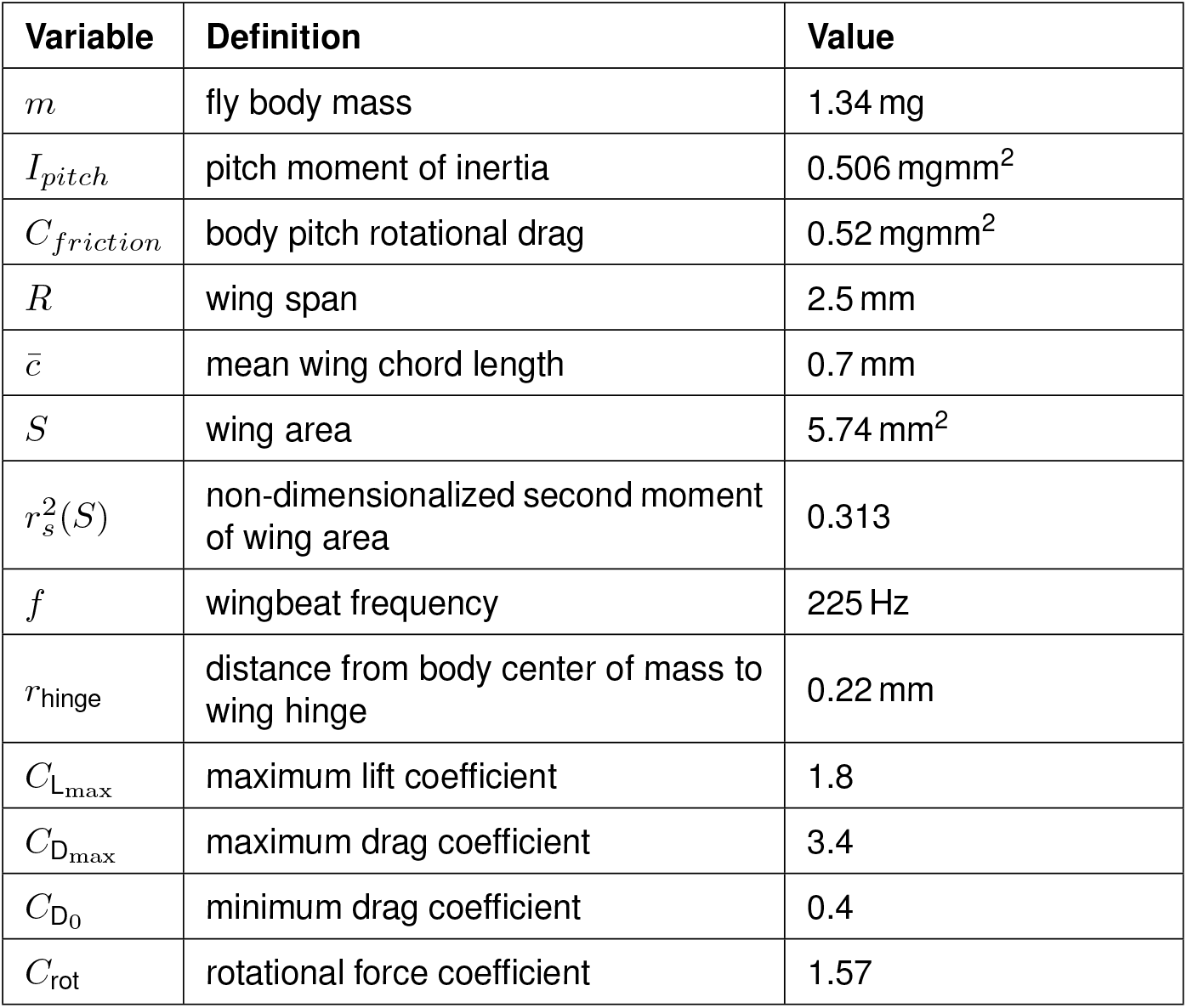
Body and wing morphological data.

## Supplementary Note 7: Full genotype of flies used in experiments

**Table 6.**
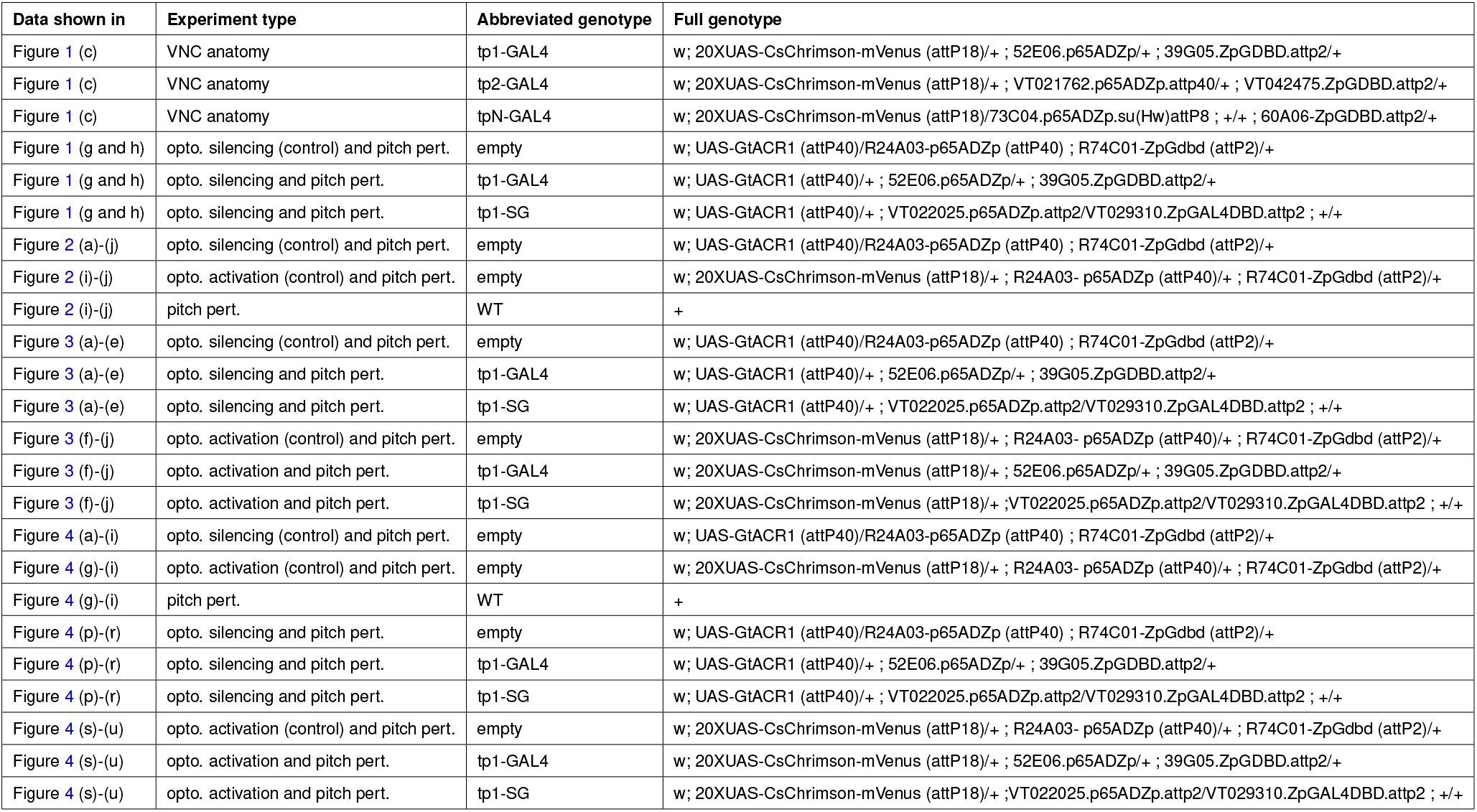
Full genotype of flies used in experiments (main text figures).

**Table 7.**
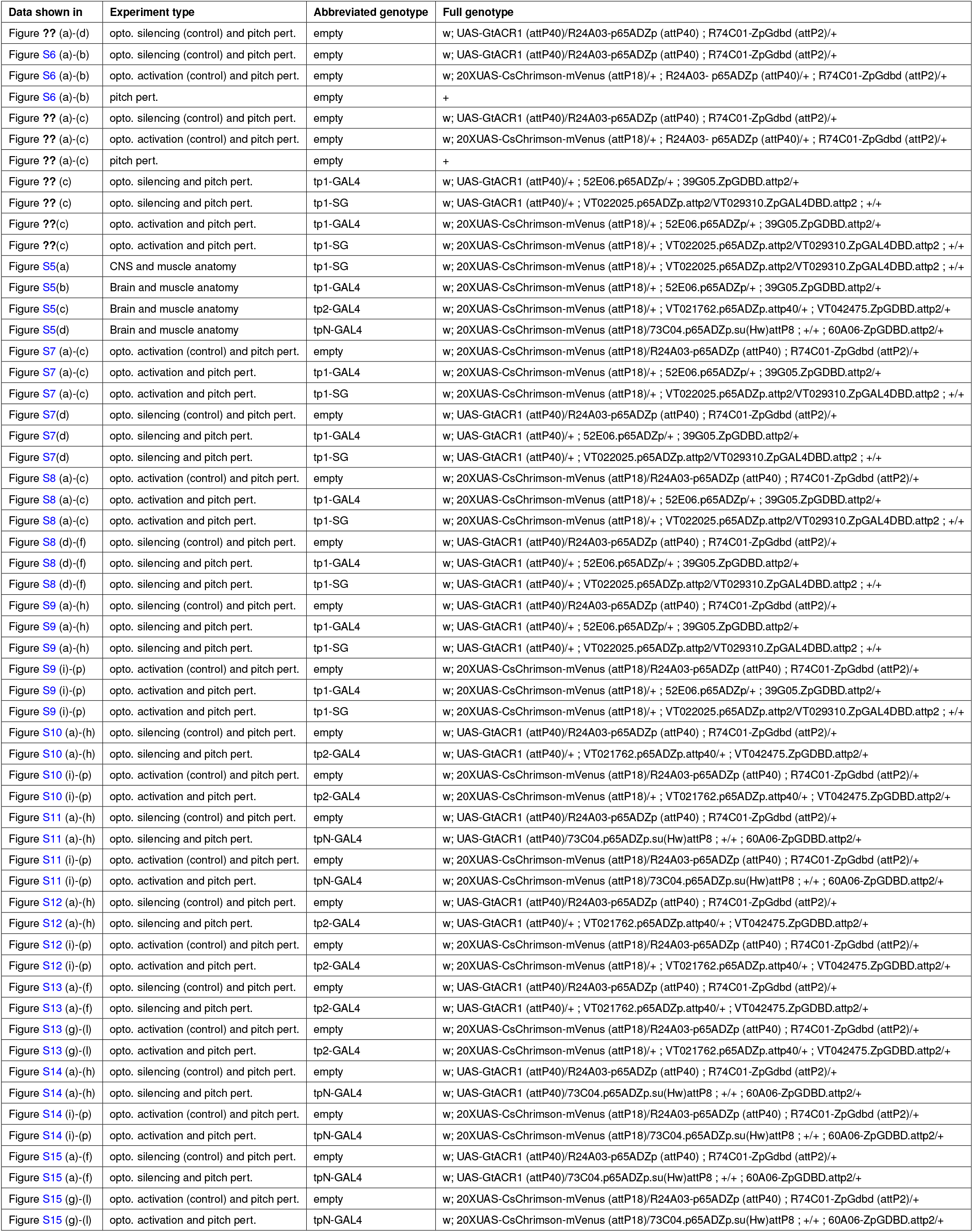
Full genotype of flies used in experiments (SI figures).

